# A genetically defined midbrain-pontine circuit gates vocal communication

**DOI:** 10.64898/2026.01.21.700949

**Authors:** Qingliang Liao, Ruiming Xu, Yang Qiao, Wenting Huang, Xian Zhang

**Author notes:** These authors contributed equally (Qingliang Liao, Ruiming Xu).

## Abstract

Vocal communication is selectively expressed in social and emotional contexts^1-3^, suggesting the presence of neural substrates that gate the conversion of internal states into vocal motor output. Although the periaqueductal gray (PAG) is known to be essential for vocalization^4-6^, the genetic identity and circuit logic of the neurons that initiate and shape vocal signals remain unclear. Here, we identify a genetically defined population of somatostatin-expressing neurons in the lateral and ventrolateral PAG (l/vlPAG^SST^) that functions as a premotor gate for ultrasonic vocalizations in mice. In vivo calcium imaging shows that l/vlPAG^SST^ neurons are selectively recruited around vocal onset. Manipulating l/vlPAG^SST^ neuron activity bidirectionally regulates vocal output: activation prolongs call duration by coordinating respiratory, laryngeal, and orofacial motor programs, whereas silencing suppresses courtship vocalizations. Projection-specific manipulations further demonstrate that descending glutamatergic, but not neuropeptidergic, output from l/vlPAG^SST^ neurons to the dorsal pontine tegmentum is sufficient to drive vocal production. Together, these findings define a genetically specified midbrain–pontine circuit that gates vocal communication and provides prospective access for linking internal conditions to vocal expressions.

## Main Text

Vocalization is among the most ancient and evolutionary conserved forms of animal communication^2,3,7-9^. From rodent ultrasonic courtship songs to human speech, vocal signals convey essential information about emotional state, motivation, individual identity, and reproductive status^1,10-14^. Generating these signals requires a hierarchical neural transformation, converting forebrain representations of internal states into precise vocal motor commands that coordinate respiration, laryngeal function, and articulation^13,15-18^. Consistent with this framework, vocalization serves as a behavioral phenotype in pathological conditions, with impaired or excessive vocal output observed in both humans and genetic animal models of neuropsychiatric disorders, including autism spectrum disorder^19-21^ and Tourette syndrome^22,23^.

At the core of this conserved network lies the midbrain periaqueductal gray (PAG), a pivotal hub that integrates emotional and sensory inputs to orchestrate vocal output by engaging downstream motor neurons^5,24-28^. Despite its central role in vocalization, the precise neuronal elements and circuit architecture within the PAG remain difficult to resolve owing to the region’s pronounced functional and cellular heterogeneity^29^. In addition to vocal control, the PAG is implicated in pain modulation, defensive behaviors, autonomic regulation, and diverse motor outputs^30-33^. As a result, bulk stimulation or inactivation of the PAG cannot readily distinguish vocalization-specific circuits from neighboring, intermingled populations. Electrophysiological recordings^34,35^ or activity-dependent tagging^4,6^ have identified PAG neurons recruited during vocal behavior, suggesting the existence of vocalization-related neural ensembles. However, whether genetically defined PAG vocal neurons exist that can be prospectively accessed to examine their causal roles in integration of forebrain inputs and initiating vocal production remains unsolved.

### Somatostatin defines a USV-activated neuronal subpopulation in the l/vlPAG

To identify PAG neurons that contribute to vocal production, we first traced descending synaptic pathways from the laryngeal musculature back into the brainstem. Injection of pseudorabies virus (PRV) into the laryngeal muscles produced time-dependent, trans-multisynaptic labeling of brainstem circuits (Extended Data Fig. 1). By 72 hours post-injection, robust PRV labeling was observed in the lateral and ventrolateral periaqueductal gray (l/vlPAG), indicating that neurons in this region are synaptically upstream of the laryngeal motor apparatus and positioned to modulate vocal output.

l/vlPAG is a longitudinal midbrain column implicated in diverse functions, including defense, itch-scratching, coughing, urination and nociception^30,36-38^. Given its anatomical linkage to laryngeal musculature, we next asked whether l/vlPAG neurons are recruited during natural vocal behavior. Most rodent species communicate using ultrasonic vocalizations (USVs) with frequencies above 20 kHz^39,40^. To reliably evoke USV production, we employed a well-established paradigm in which adult male mice were exposed to female conspecifics (Fig. 1a). Consistent with previous report^4^, Fos immunolabeling revealed a marked increase in Fos⁺ neurons within the l/vlPAG of vocalizing males, compared to non-vocalizing home-cage controls (Fig. 1b), indicating selective recruitment of this region during vocal production.

**Fig. 1.**
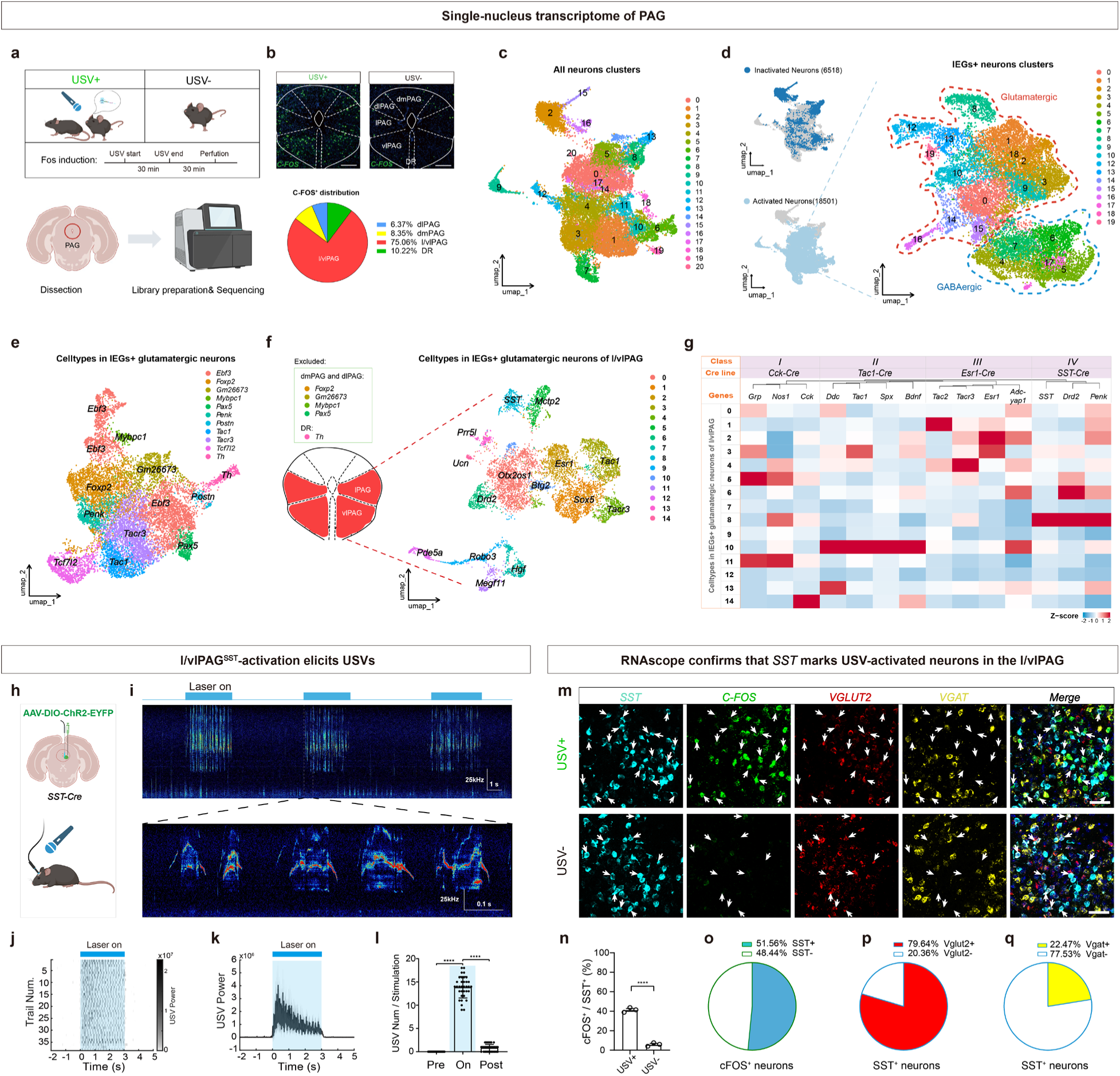
*SST* defines a vocalization-activated neuronal subpopulation in the l/vlPAG. **a,** Experimental schematic. Adult male mice were assigned to a vocalizing (USV+; courtship interaction with a female to induce Fos mRNA) or non-vocalizing (USV-; home-cage control) condition. PAG tissue punches were collected for single-nucleus RNA sequencing. **b**, Top, RNAscope detection of *Fos* mRNA in the PAG of USV+ and USV- mice. Scale bar, 200 μm. Bottom, pie chart showing the proportional distribution of *Fos*⁺ neurons across PAG subregions. **c**, UMAP embedding of all sequenced neurons (n = 25,019 nuclei from 6 mice), revealing 21 transcriptionally distinct neuronal clusters across conditions. **d,** Left, identification of activated neurons based on expression of 25 immediate-early genes (IEGs), distinguishing IEG⁺ (18,501 nuclei) from IEG⁻ (6,518 nuclei) populations across conditions. Right, UMAP embedding of re-clustered IEG⁺ neurons, yielding 20 transcriptionally defined subclusters, segregating glutamatergic and GABAergic populations. **e**, UMAP embedding of re-clustered IEG⁺ glutamatergic neurons, revealing 11 transcriptionally distinct subclusters annotated using GeneMaker. **f**, UMAP embedding of re-clustered IEG⁺ glutamatergic subclusters after exclusion of clusters preferentially localized to the dlPAG and dmPAG (*Foxp2, Pax5, Mybpc1, Gm26673*) and the dorsal raphe (*Th*), revealing 15 transcriptionally distinct subclusters enriched in the l/vlPAG. **g**, Heatmap showing scaled expression of neurotransmitter-, neuropeptide-, and receptor-related marker genes across l/vlPAG glutamatergic subclusters. Feature-based reclustering organized these neurons into four molecularly distinct classes, with corresponding available mouse Cre driver lines indicated for each class. **h**, Schematic of optogenetics activation of l/vlPAG^SST^ neurons in freely moving *SST-Cre* males following injection of AAV-EF1α-DIO-ChR2-EYFP and fiber implantation. **i**, Representative sonogram showing optogenetically evoked USVs (20 Hz stimulation; blue bar) and time-expanded view of individual syllables. **j**, Raster plot of USV syllable aligned to laser onset and (**k**) mean USV power (bottom; 2-s stimulation, 15-ms pulses; mean ± 95% CI; n = 6 mice, 38 trials), demonstrating light-evoked vocal emissions. **l**, Quantification of syllable count before, during and after optogenetic stimulation (n = 6 mice; mean ± s.e.m.; one-way ANOVA; \*\*\*\**P* < 0.0001). **m**, Representative RNAscope images comparing *SST*, *Fos*, *Vglut2*, and *Vgat* expression in l/vlPAG between USV+ and USV-mice. Arrowheads denote l/vlPAG^SST^ neurons. Scale bar, 100 μm. **n**, Percentage of *SST* neurons expressing c*Fos* under USV+ versus USV- conditions (n = 3 mice/group; unpaired t-test, \*\*\*\**P* < 0.0001). **o**, Proportion of l/vlPAG^Fos^ neurons that express *SST* in vocalizing males (n = 3). **p**, Fraction of l/vlPAG^SST^ neurons co-expressing *Vglut2* (*n* = 3 mice). **q**, Fraction of l/vlPAG^SST^ neurons co-expressing *Vgat* (*n* = 3 mice).

Genetic access to vocalization-activated PAG neurons would provide a critical entry point for circuit dissection^41^. To identify molecular markers of PAG neurons recruited during vocal behavior, we performed single-nucleus RNA sequencing (snRNA-seq) on microdissected PAG tissue from adult male mice that produced courtship USVs (USV⁺) and from non-vocalizing home-cage controls (USV⁻; n = 3 per group). Samples were processed in parallel to minimize batch effects. After quality control and data integration, 41,054 nuclei were retained, comprising eight major cell classes and 21 molecularly distinct neuronal clusters (Extended Data Fig. 2). Restricting the analysis to neurons yielded 25,019 high-quality neuronal nuclei (Fig. 1c). We next examined the expression of 25 immediate early genes (IEGs) to distinguish neurons activated during vocal behavior. This approach identified 18,501 IEG⁺ neurons across multiple clusters, compared to 6,518 IEG⁻ neurons (Fig. 1d).

Because prior studies indicate that PAG neurons engaged during ultrasonic vocalization are predominantly excitatory^4,5,42^, we focused subsequent analyses on IEG⁺ glutamatergic neurons. Re-clustering of this population yielded 11 transcriptionally distinct glutamatergic subclusters (Fig. 1e). To identify genetic entry points for functional interrogation of USV-relevant neurons within the lateral and ventrolateral PAG, we integrated transcriptional enrichment with spatial localization data from published PAG atlases^43,44^. Clusters preferentially localized to the dorsolateral or dorsomedial PAG (*Foxp2, Pax5, Mybpc1, Gm26673*) or to the dorsal raphe (*Th*) were excluded from further analysis (Fig. 1f). Re-clustering of the remaining l/vlPAG-enriched glutamatergic neurons revealed 15 transcriptionally distinct subclusters (Fig. 1f). Inspection of marker genes across these subclusters revealed considerable molecular heterogeneity, including populations defined by neuropeptides or neurotransmission-related genes (e.g., *SST, Tac1, Ucn*), receptor-associated genes (e.g., *Esr1, Drd2*), transcriptional regulators (e.g., *Sox5, Btg2*), and developmental or connectivity-related genes (e.g., *Robo3, Megf11*).

Notably, unsupervised clustering of snRNA-seq data is often driven by highly expressed transcription factors that primarily reflect developmental lineage rather than signaling features relevant for circuit function^45^. Because our objective was to identify neuron populations capable of gating vocal output, we therefore performed a targeted reclustering based on a set of neuropeptides-, neurotransmission-, and receptor-related genes derived from a prior PAG transcriptomic study^43^. Reclustering based on this gene set grouped l/vlPAG glutamatergic neurons into four molecularly distinct classes (Class *I - IV*), each with available Cre driver mouse lines (Fig. 1g), enabling systematic functional interrogation of candidate vocalization-gating populations.

We then performed an optogenetic functional screen using these genetic entry points (Extended Data Fig. 3). Channelrhodopsin-2 (ChR2) was selectively expressed in l/vlPAG neurons of *Vglut2-Cre* (General glutamatergic)*, Cck-Cre* (Class *I*)*, Tac1-Cre* (Class *II*)*, Esr1-Cre* (Class *III*)*, or SST-Cre* (Class *IV*) mice via AAV-DIO-ChR2-EYFP injection, followed by implantation of optical fibers for stimulation (Fig. 1h; Extended Data Fig. 3). Strikingly, optogenetic activation of *Vglut2⁺*, *Cck⁺*, *Tac1⁺* or *Esr1⁺* l/vlPAG neurons failed to evoke USVs. In contrast, stimulation of *SST*-expressing l/vlPAG neurons robustly and consistently elicited USVs in freely moving mice, in the absence of social or sensory cues (Fig. 1i–l; Extended Data Fig. 3; Supplementary Video 1). Notably, optogenetically evoked USVs were observed in both male and female mice (Extended Data Fig. 4), indicating that l/vlPAG^SST^ activation is sufficient to trigger vocal production independent of sex.

We next validated the anatomical and molecular identity of these neurons using RNAscope. *SST⁺* neurons were densely distributed within the l/vlPAG (Fig. 1m; Extended Data Fig. 5a, b). In vocalizing adult males, ∼44% of *SST⁺*neurons expressed *Fos*, compared to <5% in non-vocalizing controls (Fig. 1n). Conversely, *SST*⁺ neurons constituted ∼52% of all *Fos*⁺ neurons in the l/vlPAG (Fig. 1o). Dual labeling for neurotransmitter markers further showed that ∼80% of *SST⁺*neurons co-expressed the glutamatergic transporter *Vglut2*, whereas ∼20% expressed the GABAergic marker *Vgat* (Fig. 1p, q).

Mice also emit robust USVs associated with negative affective states, such as isolation^5,25^ or cold exposure^46^ in neonatal pups. To determine whether l/vlPAG^SST^ neurons are similarly engaged during these aversive forms of vocalization, we examined *SST* neuron activation in pups induced to vocalize either by removal from the home cage or by cold exposure (Extended Data Fig. 5c). In control pups maintained in the home cage, only ∼7.5% of l/vlPAG^SST^ neurons expressed *Fos* (Extended Data Fig. 5d-f). In contrast, isolation or cold exposure resulted in *Fos* expression in ∼77% and ∼81% of l/vlPAG^SST^ neurons, respectively. Conversely, *SST⁺*neurons accounted for ∼73% and ∼62% of all *Fos⁺* neurons within the l/vlPAG under isolation and cold-exposure conditions, respectively (Extended Data Fig. 5d-f), indicating that *SST* neurons comprise the dominant vocalization-activated population in this region. Dual RNAscope labeling further confirmed that the vast majority (∼91%) of l/vlPAG^SST^ neurons co-expressed the glutamatergic marker *Vglut2*, whereas only a small fraction (∼11%) expressed the GABAergic marker *Vgat* (Extended Data Fig. 5g, h).

Together, these convergent anatomical, transcriptional, and functional analyses identify l/vlPAG^SST^ neurons as a genetically defined, predominantly glutamatergic PAG subpopulation that is robustly engaged across opposing emotional contexts, from adult courtship to pup distress vocalizations.

### l/vlPAG^SST^ neuronal activity correlates with USV generation

To investigate how l/vlPAG^SST^ neurons contribute to the generation of USVs, we first monitored their bulk activity in freely moving mice during vocal behavior. Because neonatal pups are too small to permit stable optical recordings, we focused on adult courtship vocalizations as an experimentally tractable paradigm. Adult *SST-Cre* mice received l/vlPAG injections of AAV-DIO-GCaMP7s and an optical fiber was implanted for fiber photometry (Fig. 2a; Extended Data Fig. 6a). During natural courtship USV bouts, calcium signals exhibited pronounced increases that tightly coincided with USV production (Fig. 2b; Extended Data Fig. 6b, c; Supplementary Video 2). Trial-by-trial analysis revealed a reliable rise in fluorescence after USV initiation, which sustained throughout vocal emission and returned to baseline upon calling cessation (Fig. 2c, d), indicating robust recruitment of *SST* neurons during vocal output.

**Fig. 2.**
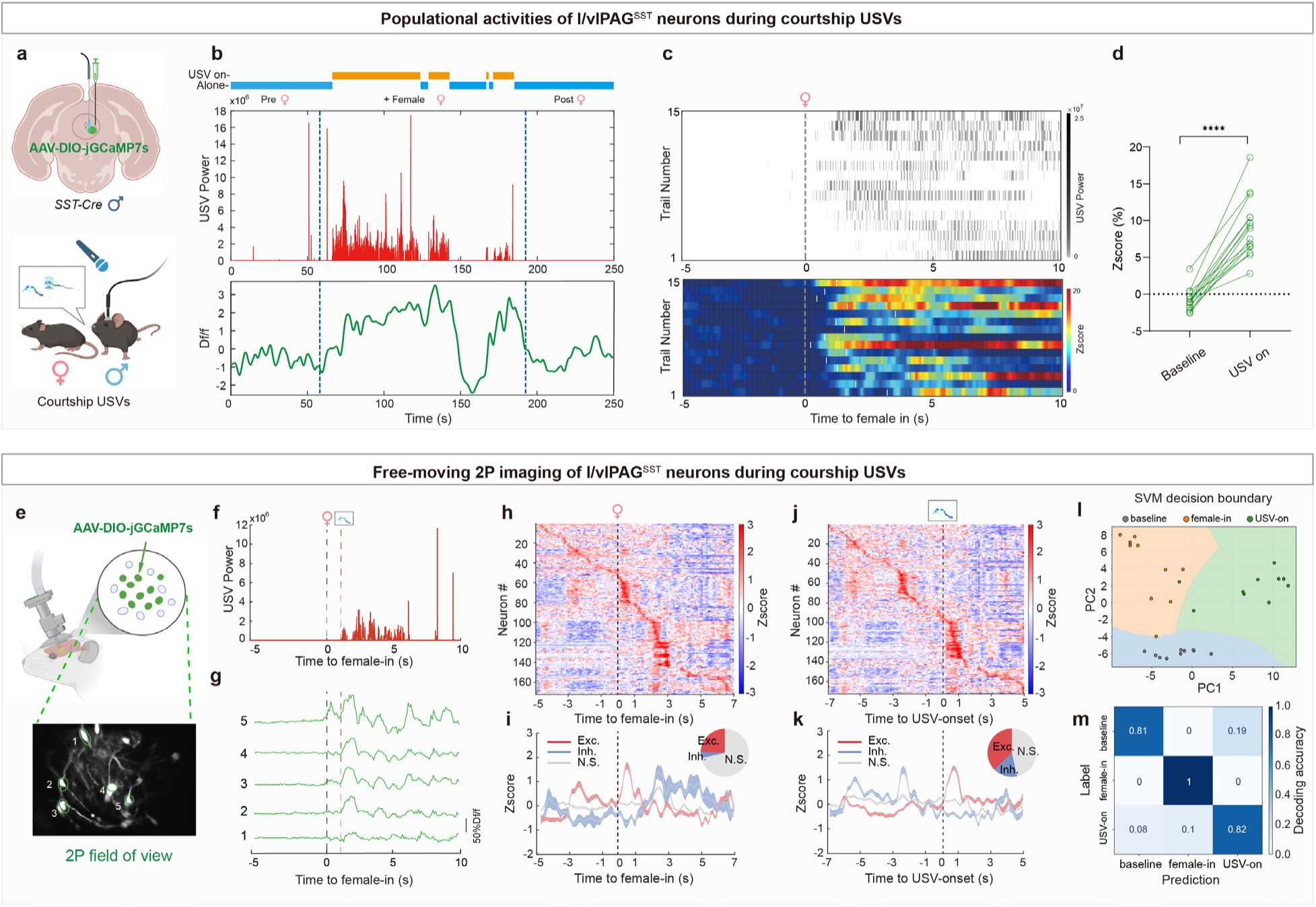
l/vlPAG^SST^ neuronal activity correlates with and predicts USV initiation during courtship. **a**, Schematics of fiber photometry recordings from l/vlPAG^SST^ neurons in male SST-Cre mice expressing AAV-DIO-jGCaMP7s during female-elicited USVs. **b**, Example trial illustrating behavioral epochs (top), USV power (middle), and corresponding calcium signals (bottom). Vertical dashed blue lines denote female introduction and removal. **c**, Trial-aligned raster plots showing USV production (top) and corresponding z-scored neural activity (bottom) from all animals. **d**, Quantification of l/vlPAG^SST^ neural activity during baseline and USV on (n = 8 mice, \*\*\*\**P* < 0.0001, paired *t*-test. **e**, Schematic of the freely moving two-photon imaging (2P) and representative field of view showing l/vlPAG^SST^ neurons expressing AAV-DIO-jGCaMP7s in male *SST-Cre* mice during female-evoked USVs. **f**, USV power trace from the same animal during imaging; black and red dashed lines indicate female entry and USV onset, respectively. **g**, Example calcium traces of individual l/vlPAG^SST^ neurons. (cell numbers correspond to panel in 2P field of view). **h**, Heat map of single-cell calcium activity from all recorded mice (n = 172 neurons from 5 mice), aligned to each neuron’s time of maximal response. Black dashed line represents the introduction of female. **i**, Mean responses of excited (red), inhibited (blue), and non-responsive (gray) neurons aligned to female introduction; shaded areas denote SEM. Inset shows the proportion of each response category (Exc.: 42 neurons, 24.4%; Inh.: 7 neurons, 4.1%; N.S.: 123 neurons, 71.5%). **j**, Heat map of single-cell calcium activity from all recorded mice (n = 172 neurons from 5 mice), aligned to each neuron’s time of maximal response. Black dashed line represents the onset of USVs. **k**, Mean responses of excited (red), inhibited (blue), and non-responsive (gray) neurons aligned to USV onset; shaded areas denote SEM. Inset shows the proportion of each response category (Exc.: 64 neurons, 37.2%; Inh.: 28 neurons, 16.3%; N.S.: 80 neurons, 46.5%). **l**, Support vector machine classifier trained on population calcium activity distinguishes baseline, female-in, and USV-on states (PC1 and PC2 projection shown). **m**, Confusion matrix showing decoding accuracy across behavioral states.

We next examined single-cell activity within this population using freely moving two-photon calcium imaging. *SST-Cre* mice expressing GCaMP7s in l/vlPAG^SST^ neurons received a gradient-index (GRIN) lens implant above the l/vlPAG (Fig. 2e; Extended Data Fig. 6d), enabling cellular-resolution imaging during courtship (Fig. 2f, g; Supplementary Video 3). We aligned the activity either to female entry or to USV onset (Fig. 2h, j), as the latency between two events varied across trials (1-2 s). Following female introduction, 24.4% of l/vlPAG^SST^ neurons exhibited increased activity (Fig. 2i). At USV onset, the proportion of activated neurons increased markedly to 37.2% (Fig. 2k). Unsupervised hierarchical clustering of USV-aligned activity revealed three functional classes (Extended Data Fig. 6e, f): Type 1 neurons that showed rapid, USV-locked excitation; Type 2 neurons that responded predominantly to female introduction; and Type 3 neurons that were largely non-responsive.

To further assess the correlation between l/vlPAG^SST^ neuronal activity and USV production, we performed state decoding using a support vector machine (SVM) classifier trained on all *SST* neuronal calcium activity during baseline, female-in, and USV-on epochs. Dimensionality reduction revealed that neural population trajectories corresponding to the three states were well segregated in low-dimensional principal-component space, with minimal overlap between conditions (Fig. 2l). When trained on these activity patterns, the classifier achieved high decoding accuracy across all states (Fig. 2m), demonstrating that population activity not only correlates with female entry and USV onset but also contains sufficient information to reliably predict the animal’s behavioral state.

Together, these population- and single-cell-level measurements demonstrate that l/vlPAG^SST^ neurons are selectively modulated upon female introduction, and are robustly recruited at the moment of vocal onset, supporting a role for this population in integrating sensory cue and gating the initiation of vocal production.

### l/vlPAG^SST^ neurons orchestrate expiratory, laryngeal, and orofacial motor programs to scale vocal output

The acoustic structure of USVs can be described by three core features: syllable duration, pitch (frequency), and loudness (amplitude)^10,47^. To determine how l/vlPAG^SST^ activity shapes these features, we optogenetically activated this population while systematically varying stimulation frequency (Fig. 3a, b; Extended Data Fig. 7; Supplementary Video 4). Stimulation at 5 Hz was sufficient to reliably elicit USVs, generating short, low-amplitude syllables. Increasing stimulation frequency progressively lengthened syllable duration across all mice, yielding a monotonic relationship between neural drive and call duration (Extended Data Fig. 7b). In contrast, USV pitch remained stable across stimulation frequencies (Extended Data Fig. 7c). USV loudness increased only at higher stimulation frequencies (20 and 40 Hz). Thus, l/vlPAG^SST^ activity scales call duration and amplitude but not pitch, indicating selective control over temporal and aerodynamic features of vocal output.

Because mammalian vocalization depends on precisely coordinated expiratory airflow and vocal fold adduction^4,16,25,48,49^, we investigated whether l/vlPAG^SST^ neurons directly control these physiology responses. Simultaneous recordings of respiration and USVs during l/vlPAG^SST^ stimulation revealed that higher stimulation frequencies produced prolonged expiratory phases, evident as flat exhalation periods in respiratory traces (Fig. 3a-e; Supplementary Video 4). The duration of flat expiration strongly correlated with the duration of the evoked USVs (R² = 0.72; Fig. 3i), indicating that l/vlPAG^SST^ neurons coordinate expiration timing with vocal output.

**Fig. 3.**
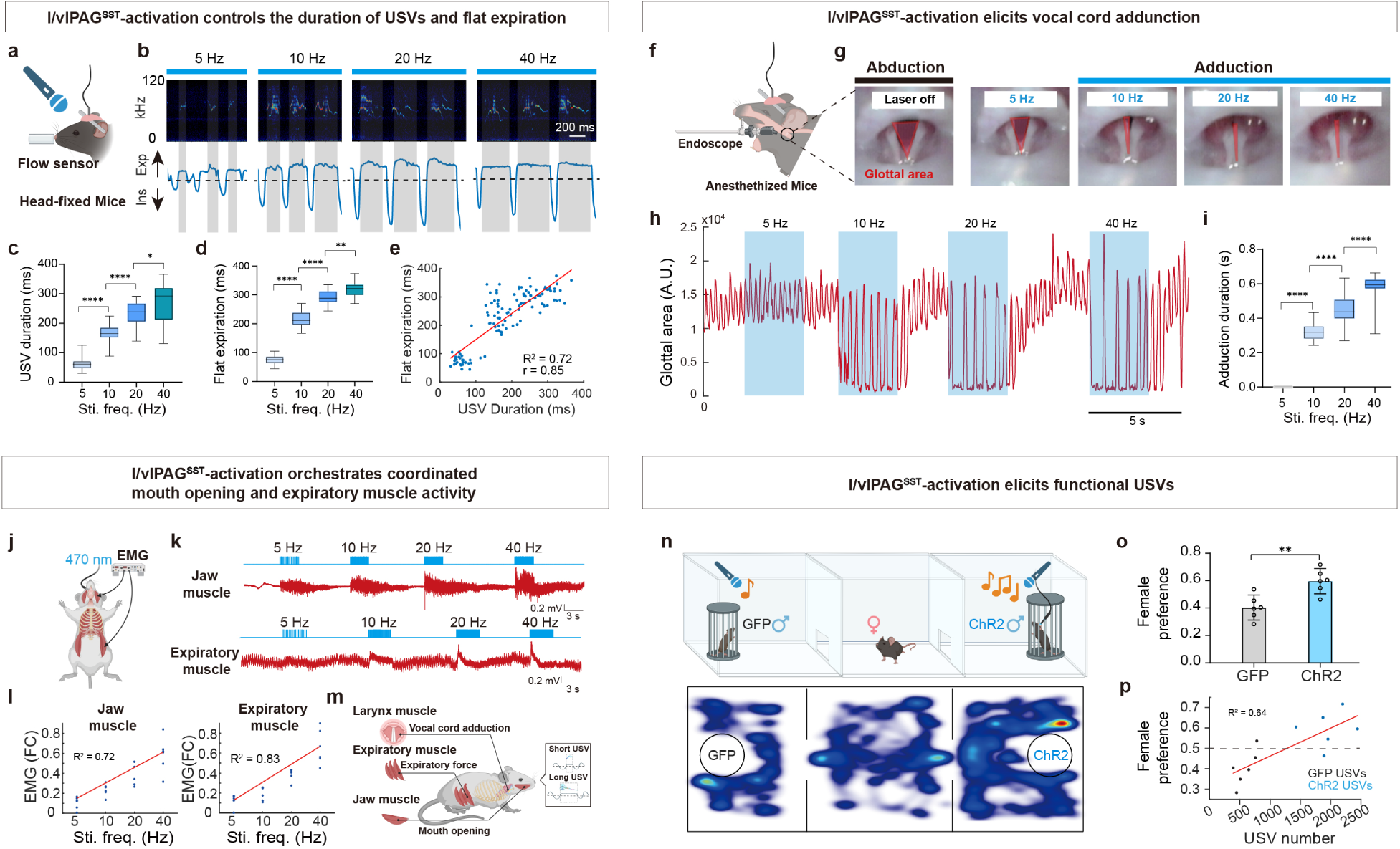
l/vlPAG^SST^ neurons coordinate expiratory, laryngeal, and orofacial motor programs to scale vocal output and promote social attraction. **a**, Schematic of simultaneous USV recording and respiratory monitoring in head-fixed mice. **b**, USV syllables (top) and respiratory traces (bottom) evoked by stimulation at increasing frequencies (5, 10, 20, and 40 Hz). Gray shading marks flat-expiration epochs. **c, d,** Frequency-dependent modulation of USV syllable duration (**c**) and flat-expiration duration (**d**) (n = 6 mice; box plots show 25th/75th percentiles, whiskers = min/max; one-way ANOVA; \*\*\*\**P* < 0.0001; \**P* < 0.05). **e**, Positive correlation between syllable duration and flat-expiration duration (linear regression; R² = 0.72; 30 trials from 6 mice). **f**, Schematic of endoscopic imaging the vocal cords in anesthetized mice. **g**, Vocal fold dynamics during baseline (abduction) and optogenetic activation (adduction) at increasing stimulation frequencies; red triangles mark the glottal area. **h**, Illustration of adduction duration evoked by stimulation at increasing frequencies (5, 10, 20, and 40 Hz). **i**, Quantification of adduction duration during laser stimulation at different stimulation frequencies (3-s epochs; n = 6 mice; box plots as in G, H; one-way ANOVA; \*\*\*\**P* < 0.0001). **j**, Schematic of EMG recording from jaw and expiratory muscles in anesthetized *SST-Cre* males with optogenetic stimulation of *SST* neurons. **k**, Representative EMG traces from jaw (top) and expiratory (bottom) muscles during stimulation at 5 - 40 Hz. **l**, Frequency-dependent increases in integrated EMG amplitude (fold change from baseline) for jaw (left; R² = 0.72) and expiratory (right; R² = 0.83) muscles (*n* = 6 mice). **m**, Model: l/vlPAG^SST^ neurons coordinate active expiration, vocal-fold adduction, and jaw opening to generate and scale USV duration. **n**, Three-chamber social preference assay testing female attraction toward control (GFP-expressing) versus vocal (ChR2-expressing) males. Bottom, representative heatmaps of female trajectory. **o**, Female preference index for GFP-expressing (gray) versus ChR2-expressing (green) males (n = 6 groups). Data are mean ± s.e.m; \*\**p* < 0.01, one-way ANOVA. **p**, Linear regression analysis showing positive correlation between female preference and the total number of male-emitted USVs (R² = 0.64; n = 6 groups).

We then examined vocal fold dynamics using endoscopic imaging in anesthetized mice (Fig. 3f). Under baseline conditions, the glottis rhythmically opened and narrowed with respiration. Optogenetic activation of l/vlPAG^SST^ neurons produced clear, frequency-dependent vocal fold adduction, and higher stimulation frequencies prolonged each addunction event (Fig. 3g-i; Supplementary Video 5).

To further probe the muscular mechanisms underlying USV production^18^, we recorded electromyograms (EMG) from two key muscles involved in vocalization during optogenetic stimulation of l/vlPAG^SST^ neurons (Fig. 3j). We targeted the masseter muscle, which controls jaw opening, and the external oblique muscle, a major abdominal expiratory muscle that generates airflow required for vocalization^25,50^. In anesthetized mice, photostimulation of l/vlPAG^SST^ neurons elicited rapid increases in EMG activity in both muscles (Fig. 3k). The magnitude of EMG responses scaled with stimulation frequency, indicating a graded recruitment of orofacial and respiratory motor output (Fig. 3l). Together with our respiratory airflow and vocal fold imaging results, these data demonstrate that activation of l/vlPAG^SST^ neurons synchronously engages expiratory muscle contraction, vocal fold adduction, and mouth opening. This coordinated pattern of motor output provides the substrate for phonation and enables precise control of USV onset and syllable duration (Fig. 3m).

We next compared optogenetically evoked syllables to the natural male vocal repertoire. Using the automated, unsupervised syllable classification tool MUPET^51^, we found that low-frequency activation produced simple, brief syllables, whereas higher frequencies generated richer and more complex syllable repertoires (Extended Data Fig. 8a-c). Correlation analyses revealed that 10-Hz-evoked syllables most closely resembled natural female-evoked courtship USVs (Extended Data Fig. 8d), suggesting that l/vlPAG^SST^ neurons can generate a graded spectrum of vocal patterns spanning the natural repertoire.

Finally, we asked whether optogenetically evoked USVs are behaviorally meaningful^4,10^. In a three-chamber social preference assay in which males were confined to opposite side chambers (Fig. 3n), females spent significantly more time investigating the ChR2-expressing males than GFP controls (Fig. 3o), which produced fewer vocalizations under the same conditions. Across animals, female preference scores strongly correlated with male USV number (Fig. 3p; R² = 0.64). Thus, l/vlPAG^SST^-evoked USVs are not only acoustically structured but also sufficient to enhance male social attraction.

### l/vlPAG^SST^ neuronal activity is required for courtship USV production

To test whether l/vlPAG^SST^ neuronal activity is necessary for the production of social USVs, we acutely silenced these neurons during ongoing courtship USVs. *SST-Cre* males received bilateral l/vlPAG injections of a Cre-dependent AAV encoding the inhibitory opsin eNpHR3.0, followed by fiber implantation for light delivery (Extended Data Fig. 9a). In freely behaving males interacting with females, continuous trains of courtship USVs and behavior were recorded simultaneously with ultrasonic audio and video (Fig. 4a). Brief illumination with yellow light (3 s, 590 nm) during sustained vocal bouts produced an immediate and reliable interruption of ongoing USVs, which promptly resumed upon light offset (Fig. 4b–e and Supplementary Video 6). Thus, l/vlPAG^SST^ neuronal activity is acutely required to maintain courtship USVs.

**Fig. 4.**
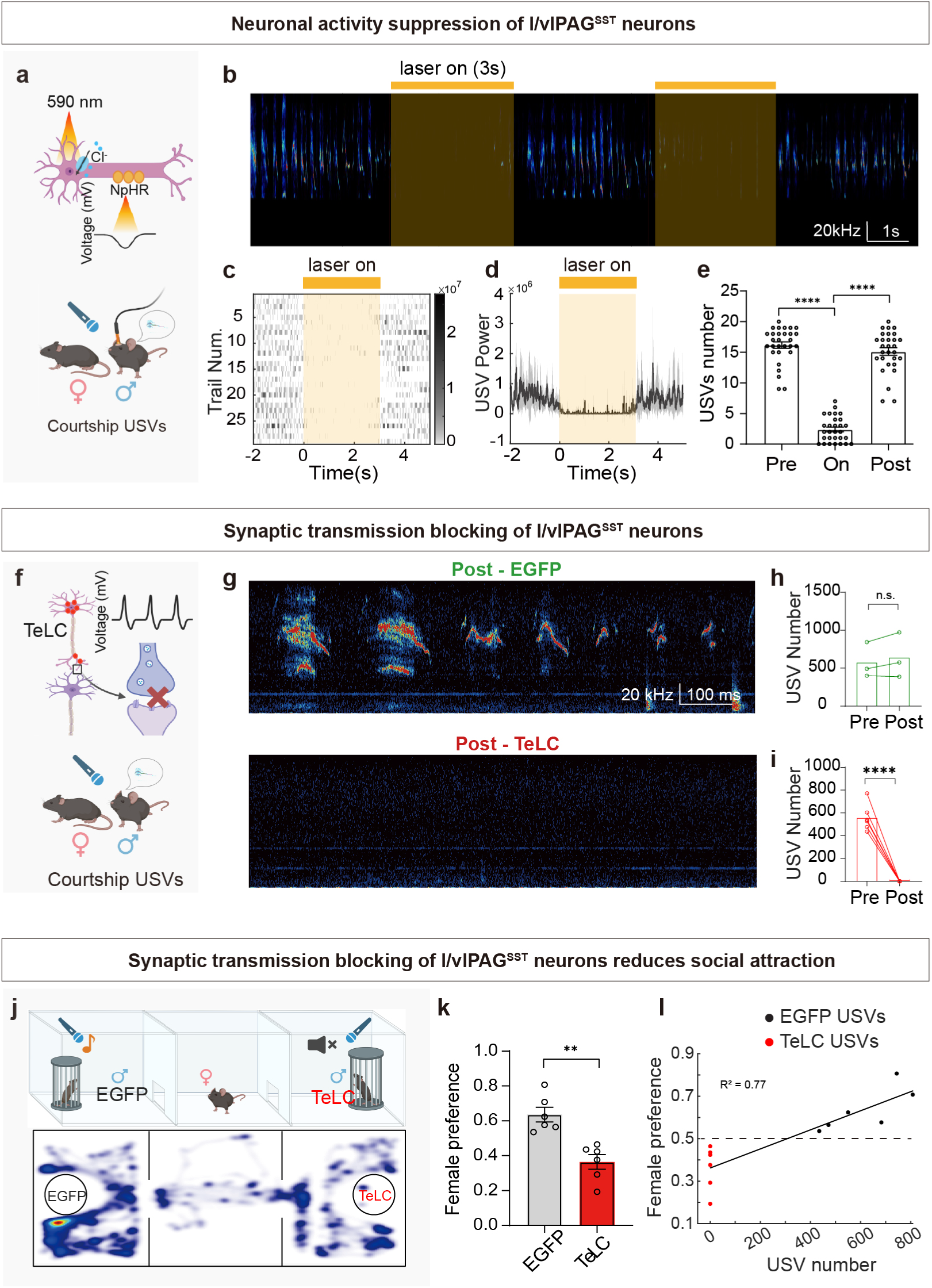
Suppressing activity or synaptic output of l/vlPAG^SST^ neurons abolishes courtship USVs and reduces social attraction. **a**, Schematic of optogenetic inhibition strategy in *SST-Cre* male mice expressing NpHR in l/vlPAG^SST^ neurons during female-elicited vocalization. **b**, Representative sonograms showing transient cessation of USVs during 3-s yellow light illumination (top bar). **c**, Trial-aligned raster plot of USV syllables demonstrating disruption of ongoing vocalizations during optogenetic inhibition (n = 6 mice; 29 trials). **d**, Mean USV power aligned to laser onset (solid line, mean; shaded region, 95% confidence interval). **e**, Quantification of USV counts before, during, and after photoinhibition (n = 6 mice; mean ± s.e.m.; one-way ANOVA; \*\*\*\**P* < 0.0001). **f**, Schematic of synaptic transmission ablation via Cre-dependent TeLC expression in l/vlPAG^SST^ neurons. **g**, Representative spectrograms showing intact USV production in EGFP controls (top) and complete loss of female-evoked USVs in TeLC-expressing animals (bottom). **h**, USV counts before and after EGFP control virus injection (n = 3; n.s.). **i**, USV counts before and after TeLC expression (n = 6; paired t-test; \*\*\*\**P* < 0.0001). **j,** Schematic of the three-chamber social preference test comparing female choice toward control (EGFP-expressing) versus muted (TeLC-expressing) males. Bottom, representative location heatmaps from one example mouse. **k**, Female preference index toward control (EGFP-expressing) males (gray) versus muted (TeLC-expressing) males (red), n = 6 groups (1 female paired with one control and one TeLC male). Data are shown as mean ± s.e.m. \*\**P* < 0.01, one-way ANOVA. **l**, Linear regression between female preference index and number of USVs emitted by males (R² = 0.77; n = 6 groups).

Because optogenetic inhibition might indirectly alter respiration or vocal motor physiology, we quantified respiratory parameters and vocal fold dynamics during inhibition. In the absence of vocalization, l/vlPAG^SST^ silencing did not significantly change breathing rate, respiratory amplitude, or baseline laryngeal movements (Extended Data Fig. 9b-g). Consistent with previous lesion studies showing that PAG damage abolishes vocalization while preserving respiration and basic laryngeal reflexes^4,52^. Together, these findings indicate that l/vlPAG^SST^ neurons function as a gate for vocal motor output, rather than as a controller of basal respiratory or laryngeal pattern generation.

We next asked whether synaptic transmission from l/vlPAG^SST^ neurons is required for USV production. To selectively block synaptic output, we expressed tetanus toxin light chain (TeLC) bilaterally in l/vlPAG of *SST-Cre* males (Fig. 4f; Extended Data Fig. 10a). Control *SST-Cre* mice expressing GFP produced robust courtship USVs both before and after viral expression (Fig. 4g, h). In contrast, all l/vlPAG^SST^-TeLC mice (n = 6) produced normal USVs prior to TeLC expression but became completely mute afterward, despite intact locomotion and courtship investigation (Fig. 4g, i; Extended Data Fig. 10b, c; Supplementary Video 7). These data demonstrate that synaptic output from l/vlPAG^SST^ neurons is essential for generating social USVs.

Given that male social USVs promote female approach^4,53,54^, we next assessed whether the loss of USVs affects social preference. In a three-chamber assay in which males were confined to opposite chambers (Fig. 4j), females spent significantly more time investigating GFP controls than TeLC-expressing muted males (Fig. 4k). Across animals, female preference scores strongly correlated with male USV production (Fig. 4l; R² = 0.77), suggesting the communicative and motivational significance of these vocal signals.

Finally, because mice also emit audible low-frequency distress calls (“squeaks”)^7,55^, we tested whether l/vlPAG^SST^ neurons are required for this non-social vocal behavior. In response to foot shock, both TeLC- and GFP-expressing males produced robust audible squeaks at comparable levels (Extended Data Fig. 10d-f). Thus, l/vlPAG^SST^ neurons are selectively required for ultrasonic social vocalizations, but dispensable for low-frequency distress calls, consistent with a functional dissociation between PAG ensembles mediating USVs versus squeaks^6^.

### l/vlPAG^SST^ neurons drive USV production via glutamatergic projections to the dorsal pontine tegmentum

To define the downstream circuits through which l/vlPAG^SST^ neurons control vocalization, we first mapped their projection targets. Following AAV-DIO-ChR2-EGFP injections into the l/vlPAG of *SST-Cre* mice (Extended Data Fig. 11a), we observed dense axonal projections to multiple forebrain and hindbrain regions, including the retroambiguus nucleus (Ram), nucleus of the solitary tract (NTS), gigantocellular reticular nucleus, dorsal pontine tegmentum (dPnTg), central amygdala (CeA), zona incerta (ZI), lateral habenula (LHb), bed nucleus of the stria terminalis (BNST), and lateral preoptic area (LPOA) (Extended Data Fig. 11b, c). Several of these structures, including LPO, NTS, Ram and ZI, have been reported to be implicated in vocal motor control^5,17,25,56,57^.

To identify which projection mediates USV generation, we selectively activated l/vlPAG^SST^ axon terminals in each downstream target. Remarkably, optogenetic stimulation of terminals in the dPnTg robustly and reliably evoked USVs, whereas stimulation of all other projection sites failed to produce vocalizations (Fig. 5a-e; Extended Data Fig. 11d); Supplementary Video 8). The stimulated region of the dPnTg encompassed Barrington’s nucleus, the locus coeruleus, adjacent pontine structures^58^, which were reported to be involved in respiration control^59^, autonomic regulation^60^, arousal^61^, urination and other motor patterning^62,63^. Consistent with this functional result, retrograde polysynaptic tracing from laryngeal muscles labeled neurons within these same pontine nuclei (Extended Data Fig. 1b), indicating that the dPnTg lies upstream of the laryngeal motor apparatus. Together, these findings identify a previously unrecognized brainstem circuit through which the PAG drives ultrasonic vocalization.

**Fig. 5.**
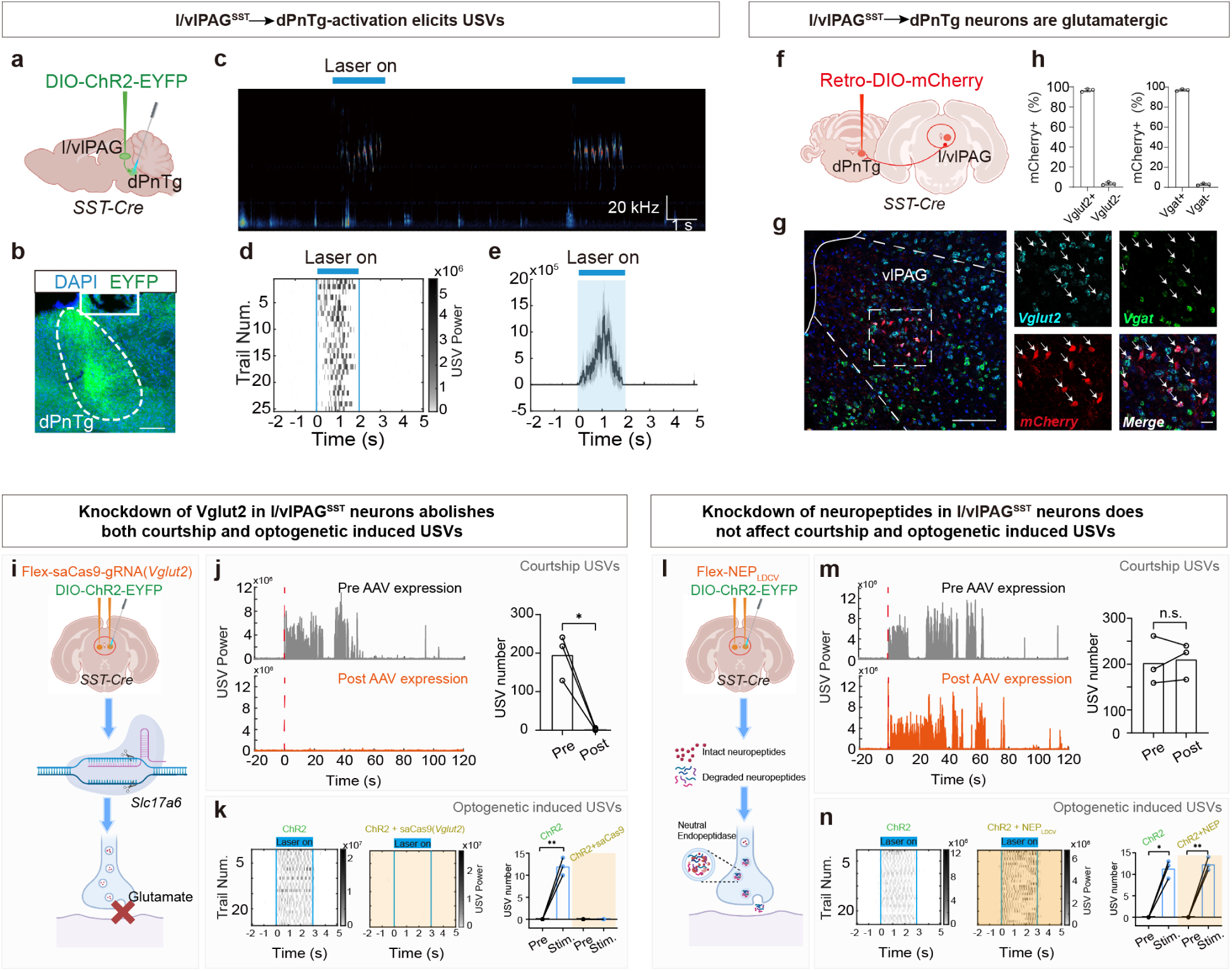
l/vlPAG^SST^ neurons drive USV production through glutamatergic projections to the dPnTg, independent of neuropeptide signaling. **a**, Schematic of AAV-DIO-ChR2-EYFP injection into l/vlPAG of *SST-Cre* mice and optical fiber implantation above dPnTg terminals to selectively activate l/vlPAG^SST^→dPnTg projections. b, ChR2-EYFP expression in dPnTg confirming targeted terminal labeling and fiber track localization. Scale bar, 200 μm. **c**, Representative sonogram showing robust USV syllables during 20-Hz photostimulation of l/vlPAG^SST^→dPnTg terminals in the absence of female interaction. **d**, Raster plot of USV syllables across trials emitted by photostimulation (25 trials, *n* = 5 mice). **e**, Mean USV power aligned to laser onset (2-s, 20 Hz), shaded region indicates 95% confidence interval (n = 5 mice, 25 trials). **f**, Retrograde AAV-DIO-mCherry labeling strategy for identifying dPnTg-projecting l/vlPAG^SST^ neurons. **g**, RNAscope images showing that mCherry-positive l/vlPAG neurons predominantly express *Vglut2* and minimally *Vgat*. Scale bars, 400 μm (left), 50 μm (right). **h**, Quantification of *Vglut2*^+^ and *Vgat*^+^ proportions among dPnTg-projecting l/vlPAG^SST^ neurons in adult males. **i**, Experimental design for *Vglut2* CRISPR knockdown and optogenetic stimulation of l/vlPAG^SST^ neurons via 1:1 co-delivery of AAV2/9-hSyn-FLEX-saCas9-HA-SV40pA-hU6-gRNA(*VGLUT2*) and together with AAV-DIO-ChR2-EYFP into l/vlPAG. **j**, Courtship-evoked USV power before and after *Vglut2* knockdown showing marked reduction in syllable number (n = 3; paired t-test, *P* < 0.05). **k**, Photostimulation-evoked USV raster plots and syllable counts demonstrating preserved vocal output in controls but complete loss following *Vglut2* knockdown (Ctrl, n = 3; *Vglut2*-saCas9, n = 3; paired t-test, *P* < 0.01). **l**, Experimental design for blocking neuropeptide signaling and optogenetic stimulation of l/vlPAG^SST^ neurons via 1:1 co-delivery of AAVDJ-DIO-NEP_LDCV_-P2A-mRuby3 and AAV-DIO-ChR2-EYFP into l/vlPAG. **m**, Courtship-evoked USVs before and after NEP_LDCV_ expression showing no significant change in syllable number (n = 3; paired t-test, n.s.). **n**, Photostimulation-evoked USV raster plots and syllable counts showing intact optogenetically induced vocalization after neuropeptide cleavage blockade (Ctrl, n = 3; NEP_LDCV_, n = 3; paired t-tests, \**P* < 0.05, \*\**P* < 0.01).

To confirm direct connectivity from l/vlPAG^SST^ neurons to the dPnTg, we performed retrograde tracing by injecting AAV2/2-Retro-DIO-mCherry into the dPnTg (Fig. 5a; Extended Data Fig. 11e). This approach selectively labeled neurons within the l/vlPAG (Extended Data Fig. 11f, g), establishing that this population sends a strong descending projection to the dPnTg. RNAscope combined with immunostaining further revealed that *SST* neurons projecting to the dPnTg are predominantly glutamatergic (∼98%, Fig. 5g, h).

Given that l/vlPAG^SST^ neurons express both the vesicular glutamate transporter *Vglut2* and the neuropeptide somatostatin, we next tested whether glutamatergic transmission, neuropeptide signaling, or both are required for USV production. To selectively disrupt glutamatergic output, we used a CRISPR-SaCas9-based strategy to knock down *Vglut2* specifically in l/vlPAG^SST^ neurons^64^. *SST-Cre* mice received bilateral l/vlPAG injections of a 1:1 mixture of AAV2/9-hSyn-FLEX-saCas9-HA-SV40pA-hU6-gRNA(*VGLUT2*) and together with AAV-DIO-ChR2-EYFP to enable optogenetic stimulation (Fig. 5i). Before viral expression, males produced robust female-evoked courtship USVs, but following *Vglut2* knockdown they became completely mute in the presence of females (Fig. 5j). Moreover, optogenetic activation of l/vlPAG^SST^ neurons failed to elicit USVs in *Vglut2*-deficient mice, whereas control males vocalized robustly (Fig. 5k). Thus, glutamatergic transmission from l/vlPAG^SST^ neurons is essential for both natural and optogenetically induced vocal output.

We next tested whether neuropeptide signaling contributes to USV generation using NEP_LDCV_, a genetically encoded enzyme that selectively degrades neuropeptides within large dense-core vesicles (Fig. 5l)^65^. Bilateral delivery a 1:1 mixture of AAVDJ-DIO-NEP_LDCV_-P2A-mRuby3 and AAV-DIO-ChR2-EYFP into the l/vlPAG led to selective inactivation of neuropeptide signaling in l/vlPAG^SST^ neurons, while allowing optogenetic stimulation of these neurons. In contrast to *Vglut2* knockdown, neuropeptide degradation had no detectable effect on female-evoked USVs (Fig. 5m) or ChR2-induced USVs (Fig. 5n), indicating that neuropeptide signaling is dispensable for USV generation.

Finally, to identify the upstream pathways that convey sensory and internal-state information to l/vlPAG^SST^ neurons, we performed monosynaptic rabies-based retrograde tracing. *SST-Cre* mice received l/vlPAG injections of Cre-dependent AAVs expressing the rabies glycoprotein (RG) and the TVA receptor fused to EGFP (Extended Data Fig. 12a). Three weeks later, EnvA-pseudotyped, glycoprotein-deleted rabies virus was injected into the same site to restrict transsynaptic labeling to monosynaptically connected presynaptic neurons. Starter cells were confined to the l/vlPAG at the injection plane (Extended Data Fig. 12b), confirming spatial specificity. Whole-brain analysis revealed a distributed but highly nonuniform input architecture. Prominent presynaptic populations were observed across multiple forebrain regions, with the densest labeling in hypothalamic, amygdala, and extended amygdala nuclei implicated in emotional state, arousal, and social behavior. Additional inputs arose from motor cortical and basal forebrain areas (Extended Data Fig. 12c-e). These results suggest that l/vlPAG^SST^ neurons constitute a major integrative hub, receiving convergent forebrain signals related to internal state and motivational context to regulate vocal output.

## Discussion

A central challenge in neuroscience is understanding how internal state and sensory context are translated into adaptive motor actions^5,14,17,66^. Vocal communication provides a powerful model for this process, yet the circuit mechanisms linking motivational signals to vocal motor output remain incompletely defined. Here we identify a genetically specified population of somatostatin-expressing neurons in the l/vlPAG that functions as a premotor gate for ultrasonic vocalizations in mice. Transcriptomic analyses and in vivo calcium imaging reveal that these neurons form a glutamatergic subpopulation selectively recruited at vocal onset, and causal manipulations demonstrate that their activity is both necessary and sufficient to initiate and scale vocal output.

Mechanistically, l/vlPAG^SST^ neurons directly engage the physiological machinery required for vocal production. Vocalizations are generated during brief, forceful expiratory events that require coordinated activation of abdominal expiratory muscles, vocal fold adduction, and mouth opening^16,18,25,49^. Optogenetic activation of l/vlPAG^SST^ neurons prolonged active expiration and increased the duration of vocal fold adduction in a frequency-dependent manner, while simultaneously recruiting jaw and expiratory musculature. This coordinated motor pattern provides a mechanistic explanation for how neural drive within the PAG is transformed into temporally scaled vocal output.

At the circuit level, we identify a specific descending pathway through the dorsal pontine tegmentum that l/vlPAG^SST^ neurons engage to drive vocal production. Given the established roles of the dPnTg in respiratory, arousal, and sensorimotor control^59,61-63^, the PAG→dPnTg pathway likely couples motivational state to premotor mechanisms underlying social vocalization. Consistent with this view, brain-wide Fos mapping recapitulated known brainstem components of the vocal network (Extended Data Fig. 13)^4,16,25^. Moreover, selective disruption of glutamatergic transmission, but not neuropeptide signaling, abolished vocal output, demonstrating that rapid excitatory signaling mediates PAG-dependent vocalization.

Taken together, by providing prospective genetic access to neurons that initiate and sustain vocal output, l/vlPAG^SST^ neurons offer a concrete entry point for linking forebrain sensory, social, and motivational signals to downstream brainstem motor programs. Our findings also provide insight into a highly conserved brainstem pathway for vocal communication, with potential translational implications for treating neuropsychiatric disorders such as autism spectrum disorder and Tourette syndrome.^19-23^.

## Methods

### Animals

*SST-Cre* mice (Jackson Laboratory, JAX 028864), *Vglut2-Cre* mice (Jackson Laboratory, JAX 039807), *Tac1-Cre* mice (Jackson Laboratory, JAX 021877), *Cck-Cre* mice (Jackson Laboratory, JAX 012706), and *Esr1-Cre* mice (Jackson Laboratory, JAX 017913) were housed and bred at the animal facility of the Hong Kong University of Science and Technology (Guangzhou). All mice were bred onto a C57BL/6J background. The animals were maintained on a 12-hour light/dark cycle with ad libitum access to food and water.

Unless otherwise noted, adult male mice (6–8 weeks old) were used for behavioral and vocalization experiments. Both sexes were included for optogenetic activation and anatomical tracing studies. All experimental procedures were approved by the Institutional Animal Care and Use Committee of the Hong Kong University of Science and Technology (Guangzhou).

### Courtship USVs induction for Fos Labeling

To evoke robust courtship ultrasonic vocalizations, adult male C57BL/6J mice (6–8 weeks old) were singly housed the day before testing to establish territorial context as previously reported. Age-matched females were group-housed and transferred onto used male bedding three days prior to testing to promote estrous cycling through pheromonal exposure. On the day of the experiment, a single female was gently placed into the resident male’s home cage. Vocal behavior was monitored in real time using an ultrasonic microphone (CM16/CMPA, Avisoft Bioacoustics) positioned above the cage to confirm the onset and persistence of USV production. Males typically produced continuous vocal bouts during the first several minutes of interaction; to standardize Fos induction across animals, interactions were allowed to proceed for 30 minutes. Immediately following the vocalization period, males were deeply anesthetized with sodium pentobarbital (100 mg/kg, intraperitoneally; MYM Technologies) and perfused transcardially with phosphate-buffered saline (PBS) followed by 4% paraformaldehyde. Brains were extracted and processed for RNAscope and immunohistochemical analyses as described below.

### Pup USVs induction for Fos Labeling

To induce distress-related ultrasonic vocalizations, neonatal C57BL/6J mouse pups at postnatal day 7 (P7) were randomly assigned to one of three experimental conditions: (1) home-cage controls, in which pups remained with their dam and littermates; (2) isolation, in which pups were individually separated from the dam and littermates for 60 min at room temperature; or (3) cold exposure, in which pups were placed individually in a temperature-controlled chamber surrounded by ice packs for 60 min. USVs were continuously recorded during the manipulation period using an ultrasonic microphone (CM16/CMPA; Avisoft Bioacoustics). Immediately following the 60-min stimulation period, pups were deeply anesthetized with isoflurane vapor and transcardially perfused for subsequent single-molecule fluorescence in situ hybridization (smFISH) analyses.

### Optogenetic activation–induced USVs for Fos labeling

To identify brain regions activated by optogenetic stimulation of l/vlPAG^SST neurons that elicit ultrasonic vocalizations (USVs), SST-Cre mice received stereotaxic injections of a Cre-dependent AAV encoding ChR2–EYFP into the l/vlPAG, followed by implantation of optical fibers above the injection site. After allowing sufficient time for viral expression and postoperative recovery, mice were individually placed in a sound-attenuating recording chamber equipped with an ultrasonic microphone (CM16/CMPA, Avisoft Bioacoustics) positioned above the cage to monitor and verify USV production. Optogenetic stimulation was delivered using blue light (10 Hz) continuously for 30 minutes to induce USVs. Sixty minutes after the end of stimulation, mice were deeply anesthetized and transcardially perfused to collect brain tissue for subsequent c-Fos immunofluorescence staining.

### Single -Nucleus RNA Sequencing

#### Experimental design and tissue collection

To investigate the transcriptional landscape of the periaqueductal gray (PAG) with social vocalization, we performed single-nucleus RNA sequencing (sn-RNAseq) on tissue from two groups: males that produced courtship USVs (n = 3) and untreated home-cage controls (n = 3). For the USV group, tissue collection occurred approximately 30 min after male–female interaction. Following euthanasia, the brains were quickly removed, and the PAG, defined by its position encircling the cerebral aqueduct, was carefully dissected. was microdissected from 1-mm coronal sections and snap-frozen in liquid nitrogen to preserve nuclear RNA.

#### Single-nucleus suspension preparation and sequencing

Frozen PAG tissues were chopped into small pieces and digested at 37°C for about 30 minutes. After digestion, the cell suspension was centrifuged, and the supernatant was discarded. Cells were resuspended with 1 mL of 1640 containing 2% FBS. After filtration through a 30-µm strainer, the suspension was adjusted to 700–1200 nuclei/µL and kept on ice. Then, cells were loaded into the microfluidic chip of Chip A Single Cell Kit v2.1(MobiDrop (Zhejiang) Co., Ltd., cat. no. S050100301) to generate droplets with MobiNova-100(MobiDrop (Zhejiang) Co., Ltd., cat. no.A1A40001). Each cell was involved in a droplet which contained a gel bead linked with up to millions of oligos (cell unique barcode). After encapsulation, droplets suffer light cut by MobiNovaSP-100(MobiDrop (Zhejiang) Co., Ltd., cat. no.A2A40001) while oligos diffuse into the reaction mix. The mRNAs were captured by cell barcodes with cDNA amplification in droplets. Following reverse transcription, cDNAs with barcodes were amplified, and a library was constructed using the High Throughput Single-Cell 3’ Transcriptome Kit v2.1 (MobiDrop (Zhejiang) Co., Ltd., cat. no. S050200301) and the 3’ Dual Index Kit (MobiDrop (Zhejiang) Co., Ltd., cat. no. S050300301)^48,49^. The resulting libraries were sequenced on an Illumina System according to the manufacturer’s instructions(Illumina) at CHI BIOTECH CO., LTD.

#### Quality control

Fastp was used to perform basic statistics on the quality of the raw sequencing reads. Raw data (fastq format) of single-cell 3’ transcriptome were pre-analyzed by MobiVision (version 3.2, MobiDrop), and reads were aligned to the reference GRCm38. A filtered cell-gene matrix was obtained with MobiVision. Specifically, for each sample, the raw (raw_feature_bc_matrix) and filtered (filtered_feature_bc_matrix) matrices were imported into SoupChannel objects. Cluster assignments were obtained from an initial Seurat object, and contamination fractions were estimated using the autoEstCont function. Corrected UMI counts were generated through the adjustCounts method, leading to the creation of new Seurat objects for each sample based on these corrected matrices. Quality control was conducted on each corrected object by calculating per-cell metrics, including the number of detected genes (nFeature_RNA), total UMI counts (nCount_RNA), and the percentage of mitochondrial transcripts (percent.mt). Cells that had fewer than 200 or more than 5,000 detected genes, or exhibited over 5% mitochondrial content, were excluded from further analysis.

#### Normalization, integration, and clustering

The Seurat package ^67^ was used for data normalization, dimensional reduction, and clustering. Datasets from different animals were integrated via canonical correlation analysis (CCA) ^68^. Highly variable genes were used for PCA, and clusters were defined at a resolution of 0.6. Differentially expressed genes (DEGs) between conditions were identified using edgeR.

#### Cluster annotation and marker gene analysis

Cluster-specific marker genes were determined utilizing the FindAllMarkers function with the MAST method. Only those genes expressed in a minimum of 10% of cells within the cluster, exhibiting a log2 fold change of at least 0.5 and an adjusted p-value of less than 0.05 (after Bonferroni correction), were included. Canonical marker genes representing major brain cell types were employed to assign cell type identities. For instance, *Snap25*, *Syp*, *Tubb3*, and *Elavl2* were identified for neurons; *Gfap*, *Aqp4*, *Slc1a3*, *Aldh1/1* for astrocytes; *Mbp*, *Plp1*, *Mog* for oligodendrocytes; *Pdgfra* and *Cspg4* for oligodendrocyte precursor cells (OPCs); *Pecam1* and *Flt1* for endothelial cells; *Tmem119*, *P2ry12*, and *Cx3cr1* for microglia; Slc17a6 and Slc17a7 for glutamatergic neurons; Gad1, Gad2, and Slc32a1 for GABAergic neurons; and Th for dopaminergic neurons. The annotation process was further supported by hierarchical clustering and the average expression patterns of these canonical markers across different clusters. Neuropeptide gene expression was assessed using manually curated gene lists, and cluster-specific expression levels were quantified (including mean log2FC, adjusted p-value, and the percentage of expressing cells) via the Wilcoxon rank-sum test implemented in the FindAllMarkers function.

#### Neuronal subclustering

The Seurat object was first filtered to retain neuronal cells, followed by unsupervised clustering. Examination of the resulting clusters revealed substantial imbalances in sample representation, with some clusters dominated by cells from a single biological replicate (≥80%), while at least one of the remaining samples contributed fewer than 10% of cells. These imbalances likely resulted from minor anatomical variability during tissue dissection, whereby adjacent regions were included in some animals but not others. To improve robustness and minimize sample-driven bias, clusters with <10% contribution from any biological sample were excluded from further analysis.

The remaining neuronal clusters, each with >10% representation from all samples, were re-clustered, yielding 21 neuronal subclusters. To identify PAG neuronal populations activated during ultrasonic vocalization (USV) production, we quantified the expression of a curated panel of 25 immediate early genes (IEGs; *Arc, Bdnf, Cdkn1a, Dnajb5, Egr1, Egr2, Egr4, Fos, Fosb, Fosl2, Homer1, Junb, Nefm, Npas4, Nptx2, Nr4a1, Nr4a2, Nr4a3, Nrn1, Ntrk2, Rheb, Scg2, Sgsm1, Syt4,* and *Vgf*) across neuronal subclusters and compared expression levels between USV and control conditions.

Neuronal populations were classified as activated if IEG expression in the USV group was significantly elevated relative to controls. Based on this criterion, neuronal clusters 0, 1, 3, 4, 5, 6, 7, 9, 11, 12, and 18 were identified as USV activated. Cells from these activated clusters were pooled and subjected to a second round of unsupervised clustering, revealing 20 transcriptionally distinct IEG⁺ neuronal subclusters.

To further refine molecularly defined entry points, glutamatergic neurons within the pooled IEG⁺ population were isolated and re-clustered, resulting in 11 transcriptionally distinct glutamatergic subclusters, which were annotated using GeneMaker. To focus specifically on lateral and ventrolateral PAG, populations relevant to vocalization control, transcriptional enrichment patterns were integrated with spatial localization information from published PAG atlases ^43,44^. Subclusters preferentially localized to the dorsolateral or dorsomedial PAG (*Foxp2, Pax5, Mybpc1, Gm26673*) or to the dorsal raphe (*Th*) were excluded. Re-clustering of the remaining l/vlPAG-enriched glutamatergic neurons yielded 15 transcriptionally distinct subclusters.

### Stereotaxic Surgery

All surgery was performed under aseptic conditions and body temperature was maintained with a heating pad. Standard surgical procedures were used for stereotaxic injection and GRIN lens or optical fiber implantation, as previously described ^69,70^. Briefly, mice were anesthetized with isoflurane (3-5% induction, 1%-1.5% maintenance, RWD Life Science) and secured in a stereotaxic frame (RWD, 71000-S), ensuring proper skull alignment relative to the bregma and lambda. Body temperature was maintained by a heating pad (37℃, RWD, ThermoStar), and ophthalmic ointment (Puralube) was applied to prevent corneal dehydration. After making a midline scalp incision to expose the skull, a microdrill (RWD, 78001) was used to create a burr hole at the coordinates corresponding to the targeted brain region.

Viral vectors were delivered using pulled glass micropipettes (tip diameter 10–30 µm) connected to an oil-driven injection system (RWD, R-480). Each site received 300–350 nL of solution at a rate of 0.3 nL/sec. Following infusion, the pipette was left in place for 10 minutes to allow diffusion and minimize backflow, then withdrawn slowly. The burr hole was sealed with bone wax, and the incision was closed with sutures. Mice were placed on a heated pad to recover before being returned to their home cages. Postoperative analgesia was provided with meloxicam (1–2 mg/kg, subcutaneous), administered immediately after surgery and continued for 2–3 days. Animals were monitored daily for discomfort or complications throughout the recovery period.

Anterior-posterior (AP) and medial-lateral (ML) coordinates are from the bregma, and dorsal-ventral (DV) coordinates are from the brain surface. The stereotaxic coordinates used in this study were as follows: l/vlPAG: AP: –4.36 mm, ML: ±1.0 mm, DV: –2.4 mm (10° ML angle) for bilateral injection, AP: –4.36 mm, ML: 0.5 mm, DV: –2.7 mm for unilateral injection. dPnTg: AP: –5.45 mm, ML: ±0.6 mm, DV: –3.5 mm; Ram: AP: –7.65 mm, ML: ±1.2 mm, DV: –5.45 mm. LPO: AP: 0.2 mm, ML: ±0.75 mm, DV: –5.2 mm; NTS: AP: –7.2 mm, ML: ±0.65 mm, DV: –4.5 mm; GiV: AP: –6.84 mm, ML: ±0.6 mm, DV: –5.75 mm.

For head-fixed experiments, mice were implanted with a custom-fabricated steel head post. For optogenetic manipulations and fiber photometry recordings, optic cannulas (200-μm core diameter, 0.37 NA; RWD Life Science) were positioned directly above the targeted brain region. For two-photon imaging, a 0.5-mm graded-index (GRIN) lens (Go!Foton) was lowered slowly into the PAG to minimize tissue deformation. A triangular head bar (Transcend Vivoscope, China), used for stabilizing the animal during miniature microscope mounting, was affixed to the skull base. All implants were secured using dental acrylic (3M Metabond) following completion of viral injections.

### Viruses

The following viral vectors were used in this study. For multisynaptic retrograde tracing from the laryngeal musculature, PRV-CAG-EGFP (Brain Case, #BC-PRV-531-plus) was injected into the intrinsic larynx muscles. For calcium imaging experiments (fiber photometry and two-photon imaging), rAAV-EF1α-DIO-jGCaMP7s-WPRE-hGH pA (BrainVTA, #PT-4552) was unilaterally delivered into the l/vlPAG of *SST-Cre* mice. For optogenetic activation, rAAV-EF1α-DIO-hChR2(H134R)-EYFP-WPRE-hGH pA (BrainVTA, #PT-0001) was injected unilaterally into the l/vlPAG. For optogenetic inhibition, rAAV-EF1α-DIO-eNpHR3.0-mCherry-WPRE-hGH pA (BrainVTA, #PT-0007) was injected bilaterally into the same site. For synaptic silencing, rAAV-EF1α-DIO-tetToxLC-P2A-mCherry-WPRE (BrainVTA, #PT-2139) was injected bilaterally into the l/vlPAG of *SST-Cre* mice. Control animals received rAAV-EF1α-DIO-EGFP-WPRE-hGH polyA (BrainVTA, #PT-1583). For monosynaptic retrograde tracing of l/vlPAG^SST^ neurons, a 1:1 mixture of AAV2/5-hEF1a-DIO-RVG (Taitool, #S0325-5) and AAV2/5-hEF1a-DIO-H2B-EGFP-T2A-TVA (Taitool, #S0320-5) was stereotaxically injected into the l/vlPAG (300 nl total). After a 3-week expression period, EnvA-SAD-B19ΔG-mRuby3 (Taitool, #R002; 200 nl) was injected into the same coordinates, and animals were perfused 7 days later. For combined Vglut2 knockdown and optogenetic activation, a 1:1 mixture of AAV-FLEX-SaCas9-HA-SV40Pa-hU6-gRNA(Vglut2) (Taitool, #S1325-9) and rAAV-EF1α-DIO-hChR2(H134R)-EYFP-WPRE-hGH pA (BrainVTA, #PT-0001) (400 nl total) was bilaterally injected into the l/vlPAG. Control animals received rAAV-EF1α-DIO-hChR2(H134R)-EYFP-WPRE-hGH pA (BrainVTA, #PT-0001) injection. For combined neuropeptide degradation and optogenetic activation, *SST-Cre* mice were bilaterally injected with a 1:1 mixture of AAVDJ-DIO-NEPLDCV-P2A-mRuby3 and rAAV-EF1α-DIO-hChR2(H134R)-EYFP-WPRE-hGH pA (BrainVTA, #PT-0001) (400 nl total). Control mice received rAAV-EF1α-DIO-hChR2(H134R)-EYFP-WPRE-hGH pA (BrainVTA, #PT-0001) injection. To validate the l/vlPAG → dPnTg descending projection, rAAV-EF1α-DIO-mCherry-WPRE-hGH pA Retro (BrainVTA, #PT-0013) was injected unilaterally into the dPnTg of *SST-Cre* mice. All viruses were aliquoted and stored at –80 °C until use.

### Retrograde Trans-Multisynaptic Tracing from Laryngeal Muscles

To identify brainstem regions involved in the control of laryngeal muscles, PRV-based peripheral retrograde tracing was performed. Wild-type mice were anesthetized with isoflurane (3% induction, 1.5% maintenance), and a midline incision was made in the ventral neck region. The sternohyoid muscles were gently dissected and retracted bilaterally to expose the larynx. PRV-CAG-EGFP was injected into the laryngeal muscles (2.5 µL per site) using a quartz micropipette (Hamilton) connected to a programmable syringe pump (KDS Legato 130). Animals were perfused for histological analysis at 24, 48, or 72 h following injection.

### Retrograde monosynaptic tracing with pseudotyped rabies virus

To map the brain-wide monosynaptic inputs onto l/vlPAG^SST^ neurons, we used cell-specific tracing strategy with an optimized rabies virus system as previously described. Briefly, we first injected the l/vlPAG of *SST-Cre* mice with AAV-DIO-RVG and AAV-DIO-HEB-EGFP-T2A-TVA, which allows Glycoprotein and TVA to be specifically expressed in l/vlPAG^SST^ neurons flowing. Three weeks after the first injection, mice were injected in l/vlPAG with ENVA-SAD-B19ΔG-mRuby3, a rabies virus that is pseudotyped with EnvA, lacks the envelope glycoprotein, and expresses mRuby3. Brain tissue was prepared one week after the rabies virus injection for histological examination. This method ensures that the rabies virus exclusively infects the cells that express TVA. Furthermore, complementation of the modified rabies virus with the envelope glycoprotein in the TVA-expressing cells allows the generation of infectious particles, which can then trans-mono-synaptically infect presynaptic neurons.

We followed a commonly used method to analyze the rabies tracing data. Briefly, 25-µm coronal sections spanning the entire anteroposterior extent of the brain using a cryostat (Epredia NX70 HOMVP) were collected, and every third section was analyzed. mRuby3-labeled input neurons were manually counted and assigned to brain regions based on anatomical landmarks defined in the Allen Brain Atlas.

### Immunohistochemistry

Immunohistochemistry experiments were performed following standard procedures. Briefly, mice were anesthetized with pentobarbital sodium (100 mg/kg, i.p.) and subsequently perfused with ice-cold 1× phosphate-buffered saline (PBS), followed by 4% paraformaldehyde overnight followed by cryoprotection in a 30% PBS-buffered sucrose solution for 36 h at 4 °C. Tissue sections were permeabilized using PBS supplemented with 0.3% Triton X-100. Primary antibodies were then applied in the blocking buffer and incubated overnight at 4°C. After three washes with PBS, the sections were incubated with secondary antibodies in the blocking buffer for 1 hour at room temperature, followed by three additional washes. Finally, a coverslip was mounted using Fluoroshield Reagent (Abcam, # ab104139) to preserve the samples. Primary antibodies included: rabbit anti-cFos (Genetex, # GTX129846), rabbit anti-mCherry (Genetex, # GTX128508), and species-specific goat secondary antibodies conjugated to Alexa Fluor 594 were obtained from ABclonal and used at a 1:500 dilution.

### Fluorescent in situ hybridization

Single molecule fluorescent *in situ* hybridization (smFISH) (RNAscope, ACDBio) was used to detect the expression of *SST*, *Fos*, *Slc17a6* and *Slc32a1* mRNAs. For tissue preparation, mice were first anesthetized under isoflurane and then decapitated. Their brain tissue was first embedded in cryomolds filled with M-1 Embedding Matrix then quickly fresh-frozen on dry ice. The tissue was stored at -80 °C until it was sectioned with a cryostat. Cryostat-cut sections (20-μm) containing the entire PAG were collected rostro-caudally, and quickly stored at -80 °C until processed. Hybridization was carried out using the RNAscope kit (ACDBio).

The day of the experiment, frozen sections were post-fixed in 4% PFA in RNA-free PBS (hereafter referred to as PBS) at room temperature (RT) for 15 min, then washed in PBS, dehydrated using increasing concentrations of ethanol in water (50%, once; 70%, once; 100%, twice; 5 min each). Sections were then dried at RT and incubated with Protease IV for 30 min at RT. Sections were washed in PBS three times (5 min each) at RT, then hybridized. Hybridization was carried out for 2 h at 40 °C. After that, sections were washed twice in PBS (2 min each) at RT, then incubated with the amplification reagents for three consecutive rounds (30 min, 15 min and 30 min, at 40 °C). After each amplification step, sections were washed twice in PBS (2 min each) at RT. Finally, fluorescence detection was carried out for 15 min at 40 °C. Sections were then washed twice in PBS, incubated with DAPI for 2 min, washed twice in PBS (2 min each), then mounted with coverslip using mounting medium. Images were acquired using a full-spectrum high-resolution fluorescence lifetime laser confocal microscope (Leica, STELLARIS 8) equipped with 20x lenses, and visualized and processed using ImageJ and Adobe Illustrator.

### In vivo fiber photometry and data analysis

Population-level activity of l/vlPAG^SST^ neurons was recorded *in vivo* in freely behaving male mice using a fiber photometry system (ThinkerTech, Nanjing). Mice were allowed to recover for at least three weeks following surgery before behavioral testing. Calcium-dependent fluorescence signals were acquired using excitation at 470 nm, together with an isosbestic control signal at 405 nm to account for motion-related and non–calcium-dependent fluctuations. The two excitation wavelengths were alternated at 50 Hz, as described previously.

To ensure precise temporal correspondence across recording modalities, video acquisition and fiber photometry signals were synchronized using an analogue TTL trigger. Behavioral events, including female introduction and USV onset, as well as calcium transients, were identified and timestamped post hoc. GCaMP fluorescence signals were further aligned with USV recordings using the same synchronization framework employed in the USV analysis pipeline, integrating timing information derived from both audio and video streams. This approach enabled accurate alignment of neural activity with behavioral and vocal events at sub-second resolution.

Each recording session began with a one-minute baseline period in the male mouse’s home cage. A live female mouse was then introduced into the cage for two minutes to induce courtship-associated USVs, after which the female was removed. Fluorescence data were processed using custom MATLAB (MathWorks, Natick, MA) scripts. Relative fluorescence changes (ΔF/F₀) were calculated as (F_raw(t) − F₀)/F₀, where F₀ represents the mean raw fluorescence during the baseline period. Z-scored fluorescence signals were also computed for population-level analyses.

### Miniature microscopic two-photon imaging and data analysis

To monitor l/vlPAG^SST^ neuronal activity at single-cell resolution in freely behaving mice, we employed a miniature two-photon imaging system (Transcend Vivoscope, China). Three to four weeks after the virus injection, mice were briefly head-fixed via the implanted head bar to assess GCaMP7s expression. Once robust expression was confirmed, animals were anesthetized with isoflurane (1.5%; RWD, China), and a microscope baseplate (Transcend Vivoscope, China) was secured to the cranial implant using a blue-light–curable dental adhesive (Vertise Flow, Kerr). The microscope was then removed, the exposed GRIN lens was protected with silicone elastomer (Kwik-Cast, WPI, USA), and mice were returned to their home cages for recovery.

Before behavioral testing, the microscope was mounted onto the baseplate via head fixation. Mice were allowed to freely explore their home cages for 5 minutes of habituation on two consecutive days prior to recording. Imaging data were acquired using dedicated software (GINKGO-MTPM, Transcend Vivoscope, China) at a frame rate of 10 Hz and a spatial resolution of 512 × 512 pixels. Excitation was provided by a femtosecond fiber laser (TVS-FL-01, Transcend Vivoscope, China), delivering approximately 35 mW at the objective and yielding a field of view of 420 × 420 μm. Imaging frame timestamps and behavioral signals were synchronized via the system controller (TVS-MMM-01, Transcend Vivoscope, China).

Raw imaging stacks were processed using Suite2P ^71^ for motion correction, region-of-interest (ROI) detection, and fluorescence trace extraction, using default parameters specified in the provided ops file. Somatic ROIs were manually curated based on Suite2P-generated mean images, correlation maps, and maximum projection views to exclude non-somatic structures. Output data were further analyzed using custom MATLAB scripts. Neuropil contamination was subtracted, and frames exhibiting excessive motion artifacts were excluded based on post-registration x–y displacement metrics. Specifically, frames were removed if their phase correlation with the reference image fell below 50% of the maximum peak phase correlation observed within the imaging stack.

To determine whether a neuron was significantly (P < 0.05) excited or inhibited by a behavioral event, that is female introduction or USV onset, and thus can be classified as being “responsive” to the event, we compared fluorescent values within a 1-s baseline window preceding the event to those within a 1-s window immediately following event onset using a Wilcoxon signed-rank test (P < 0.05). Neurons showing significant increases or decreases were classified as excited or inhibited, respectively. Temporal z-scores for each neuron were calculated by first averaging ΔF/F₀ traces across trials at each time point and then normalizing these values to the baseline mean and standard deviation.

To classify neurons according to their response dynamics, principal component analysis (PCA) was performed on the z-score–normalized activity profiles as previously reported. Hierarchical clustering was subsequently applied to the first three principal components using “*correlation”* distance and “*complete”* linkage, implemented with custom MATLAB scripts.

### Decoding analysis

To test whether transient behavioral states (baseline, female introduction, and USV onset) could be predicted from population-level l/vlPAG^SST^ activity, we trained a support vector machine (SVM) classifier with a radial basis function (RBF) kernel using the activity of all recorded neurons. Because the interval between female introduction and USV onset typically ranged from 1–2 s, we restricted each condition to a 1-s window to ensure balanced class sizes, yielding three equally represented conditions. For each trial, neural population activity was represented as a 172-dimensional feature vector, with each dimension corresponding to one neuron. Prior to classification, feature vectors were z-scored and reduced using principal component analysis (PCA), and the first two principal components were used for subsequent decoding.

The SVM classifier (C = 10, γ = “scale”) was trained and evaluated within this two-dimensional PCA space. Decoding performance was assessed using stratified 10-fold cross-validation. Specifically, the dataset (30 frames in total) was randomly divided into ten equally sized folds, each containing one trial from each behavioral condition. In each iteration, nine folds were used for training and the remaining fold for testing, such that each fold served once as the test set. To reduce potential bias introduced by a particular data partition, the entire 10-fold cross-validation procedure was repeated 1,000 times with different random seeds. Decoding accuracy is reported as the mean ± s.d. across these repetitions.

To obtain a robust visualization of decision boundaries, we trained a total of 10,000 SVM classifiers (1,000 repetitions × 10 folds). For each point on a coarse grid in PC1–PC2 space (step size = 0.1), class identity was assigned by majority vote across all classifiers. The resulting classification map was then interpolated onto a finer grid (step size = 0.05) using nearest-neighbor interpolation for display. Confusion matrices are shown for a representative 10-fold cross-validation as well as for the normalized matrix averaged across all 1,000 repetitions. All decoding analyses were performed in Python 3.9 using scikit-learn (v1.3) and matplotlib (v3.8).

### USV analysis

Ultrasonic vocalizations (USVs) were analyzed using custom MATLAB scripts adapted from established Stowerslab pipelines in combination with Avisoft SASLab Pro software (Avisoft Acoustics). For each recording, raw acoustic signals were first band-pass filtered between 40 and 110 kHz to remove low-frequency environmental noise and high-frequency artifacts unrelated to mouse USVs. Time-frequency representations were then computed using the MATLAB “*spectrogram”* function, allowing estimation of the acoustic power spectrum from recordings obtained with an ultrasonic microphone (Avisoft Bioacoustics, CM16/CMPA). Spectral power within the 40–110 kHz band was converted to decibel (dB) units using a reference power of 10 × 10⁻¹² W. To further suppress background noise, power values that did not exceed 1.5 standard deviations above the mean room noise level were subtracted. The resulting power trace was subsequently smoothed with a 50-ms Gaussian kernel to reduce high-frequency fluctuations. Total USV power was quantified by integrating the smoothed power signal over time using the MATLAB “*trapz”* function. Individual USV events were detected by applying the *“findpeaks”* function to the smoothed power trace, with detection thresholds set to a minimum peak height of one standard deviation above the noise level and a minimum inter-peak interval of 150 ms. For population-level analyses, average USV power traces were calculated across trials, and variability was represented as the 95% confidence interval of the mean, computed as 1.96 × the standard error of the mean (s.e.m.).

### MUPET analysis

USV features were analyzed in MATLAB (MathWorks) using the Mouse Ultrasonic Profile Extraction Tool (MUPET) as previously reported ^51^. Audio files were processed with default MUPET parameters to segment individual ultrasonic syllables. All extracted syllables were subsequently inspected by a blinded experimenter, and segments corresponding to non-vocal artifacts (e.g., audible movement-related noise) were manually excluded from further analysis. Quantitative features of each syllable, including amplitude, duration, and peak frequency, were obtained directly from the MUPET output. For repertoire construction and comparison, 60 syllables were empirically selected to represent the vocal output in each condition. To assess similarity between optogenetically evoked vocalizations and female-induced courtship USVs, Pearson’s correlation analysis was performed on the syllable repertoires generated by MUPET.

### *In vivo* optogenetic activation and inhibition

For ChR2-mediated photostimulation experiments, fiber-implanted mice were gently handled during manual attachment and detachment of optical patch cables to minimize stress. Optogenetic activation of l/vlPAG neurons or downstream projections was achieved using a 473-nm laser (output power: 10 mW; stimulation frequency: 5–40 Hz) delivered through an optogenetic stimulation system (ThinkerTech Bioscience Co., Nanjing). Light output at the fiber tip was measured and verified both before and after each recording session using a calibrated photodiode power meter (Thorlabs PM20A). Behavioral experiments were conducted in a sound-attenuated recording chamber equipped with an ultrasonic microphone synchronized to overhead CCD cameras capturing video at 30 frames per second, following established experimental procedures.

For eNpHR-mediated optogenetic inhibition experiments, courtship-directed USV assays were performed using established paradigms. Male mice expressing eNpHR in l/vlPAG^SST^ neurons were individually housed 24 hours prior to behavioral testing. Female mice were group-housed (five per cage) and exposed to bedding soiled by male mice for 72 hours before experiments to promote estrous synchronization. On the day of testing, fiber-implanted males were gently handled during patch cable connection and allowed to habituate in the recording chamber for 15 minutes. A sexually receptive female was then introduced into the arena to reliably elicit sustained courtship USV production. Bilateral photoinhibition of l/vlPAG^SST^ neurons was achieved using 590-nm laser illumination (output power: 10 mW; pulse duration: 3 s) delivered through an optogenetic inhibition system (ThinkerTech Bioscience Co., Nanjing), with light applied during periods of active vocalization.

### Synaptic transmission inhibition

To selectively block synaptic output from l/vlPAG^SST^ neurons, we expressed tetanus toxin light chain (TeLC) in SST neurons and compared female-elicited USV production within the same male mice before and after TeLC expression.

Pre-inhibition: Male *SST-Cre* mice (2-3 months old) underwent 24-hour isolation housing before USV recording. During USV recording sessions, subjects were transferred to standardized white polycarbonate observation chambers (30×30×35 cm) equipped with ceiling-mounted Avisoft CM16/CMPA ultrasonic transducers (sampling rate: 250 kHz) synchronized with overhead CCD video capture systems (30 fps). Following a 15-minute habituation phase, stimulus females (estrus-confirmed via vaginal cytology37) were placed in the male’s cage for 3 min, and the encounter was recorded. The next day, male *SST-Cre* mice were randomized to ablation or control groups, receiving bilateral l/vlPAG microinjections of either: rAAV-EF1α-DIO-tettoxlc-P2A-mCherry-WPRES (TeLC group) or rAAV-EF1α-DIO-EGFP-WPRE-hGH polyA (EGFP group).

Post-inhibition: Following a 4-week expression period ensuring complete toxin-mediated neuronal silencing, vocalization was recorded as above in a 3-min encounter with a wild-type female in estrus. Vocalizations were assumed to be produced by males because males produce the majority of USVs during courtship behaviors. To calculate social interaction time, videos were manually scored by counting the number of seconds during the 3-minute trial in which the male mouse’s nose or forelimb was in contact with the female.

### Pain-induced audible squeak

To assess pain-evoked low-frequency vocalization, mice were subjected to brief electrical foot shocks (0.5 mA, <2 s) delivered in a standard shock chamber. Behavioral reactions and vocalizations were simultaneously recorded using an overhead video camera (30 frames per second) synchronized with an ultrasonic microphone. Audible squeaks were identified and analyzed for acoustic features, including peak frequency (kHz) and sound intensity (dB). Vocalization timing was aligned with video recordings to correlate squeak emission with concurrent behavioral responses, such as escape movements or freezing behavior.

### Three-chambered social preference test

Social preference was evaluated using a standard three-chamber apparatus consisting of a central compartment flanked by two identical lateral chambers. Female mice were initially allowed to freely explore all three chambers for a 5-min habituation period to acclimate to the apparatus. For the test phase, stimulus males were placed individually under wire mesh cages positioned in each lateral chamber, allowing transmission of olfactory and auditory cues while preventing direct physical contact. Two experimental comparisons were conducted: (1) *SST-Cre* male mice expressing EGFP (control) versus SST-Cre males expressing ChR2 and receiving continuous 10-Hz optogenetic stimulation to sustain USV production; and (2) *SST-Cre* males expressing EGFP that produced normal courtship USVs versus *SST-Cre* males expressing TeLC, which were rendered mute.

During setup, the female subject was confined to the central chamber and subsequently allowed unrestricted exploration of the entire apparatus for a 15-min test session. Social interaction time was defined as the cumulative duration the female spent within 3 cm of each stimulus cage and was quantified using EthoVision XT software (v15.0; Noldus) based on overhead video recordings acquired at 30 frames per second. Ultrasonic vocalizations emitted by stimulus males were simultaneously recorded using microphones positioned above each lateral chamber (Avisoft CM16/CMPA; sampling rate, 250 kHz). Spectrotemporal features of USVs were analyzed in parallel with female approach behavior. A preference index was calculated for each animal as (time_target − time_control)/total_time. Preference indices were statistically compared against chance levels using a one-sample t-test.

### Respiratory activity recording and analysis

Respiratory dynamics were monitored in head-fixed l/vlPAG^SST^-ChR2 or l/vlPAG^SST^-eNpHR mice using a miniature airflow sensor (Honeywell AMW3300V) positioned immediately in front of the snout. Analog voltage signals were acquired at 250 kHz using a high-speed data acquisition system (National Instruments PCIe-6321) and subsequently downsampled to 1 kHz for offline analysis. Raw airflow signals were normalized by subtracting baseline no-flow reference values and scaling to the standard deviation of resting-state respiration.

Periods of flat expiratory airflow were identified automatically using custom-written algorithms implemented in MATLAB. The timing of optogenetic stimulation, USV recordings, and respiratory signals was synchronized through an open-source Bpod-controlled interface (Sanworks, Stony Brook, NY, USA). This configuration enabled precise temporal alignment across laser delivery, ultrasonic microphone acquisition (>50 kHz sampling rate), and overhead CCD video recording at 30 frames per second. The integrated system provided sub-millisecond temporal precision across all data streams throughout behavioral sessions.

To quantify the relationship between respiration and vocal output, a linear regression model was fitted to relate the duration of USVs to the duration of flat expiratory phases. Model performance was assessed by calculating the coefficient of determination (R²).

### Vocal cord imaging and analysis

To visualize laryngeal dynamics during optogenetic manipulation of l/vlPAG^SST^ neurons, mice were deeply anesthetized by intraperitoneal injection of pentobarbital sodium (100 mg/kg) before surgical preparation. Animals were secured on a temperature-controlled platform, and the head was stabilized using custom clamps. Proper alignment of the upper airway was achieved by supporting the ventral neck to straighten the orotracheal axis, while the tongue was gently extended and immobilized under direct visualization using a flat-tipped titanium depressor.

Vocal fold movements were imaged transorally using an endoscopic system operating at 30 frames per second. Endoscopic video acquisition was synchronized with bilateral optogenetic stimulation or inhibition. For optogenetic activation, 473 nm laser pulses were delivered for 3 s at stimulation frequencies of 5, 10, 20, or 40 Hz. For optogenetic inhibition, 590 nm laser pulses were applied for 2 s using parameters consistent with those described above. Laser timing was controlled through the Bpod interface to ensure precise temporal coordination with imaging.

Quantitative analysis of vocal fold kinematics was performed using DeepLabCut-based pose estimation. Custom-trained neural networks were used to track frame-by-frame changes in the interarytenoid distance, providing a continuous measure of glottal opening and closure dynamics. Multimodal synchronization across laser triggers, endoscopic video frames, and behavioral timestamps was maintained with sub-millisecond precision (±1 ms), enabling phase-resolved analyses of vocal fold movements in relation to optogenetic stimulation.

### Masseter and expiratory muscle EMG recording

For masseter and expiratory muscle EMG recording during optogenetic stimulation of l/vlPAG^SST^ neurons, mice were deeply anesthetized via intraperitoneal administration of pentobarbital sodium (100 mg/kg) prior to surgical preparation. The mouse was positioned supine, and the skin over the expiratory muscle was aseptically prepared. A 1-cm incision was made to expose the expiratory muscle, after which a two-lead needle electrode (ADInstruments, MLA1203) was carefully inserted into the muscle. The masseter muscle was similarly exposed through a 1-cm incision made under the chin, followed by careful dissection of the overlying strap muscles. Electrodes were connected to an ADInstruments Octal Bio Amp and an ADInstruments PowerLab data acquisition system, which recorded EMG signals at a sampling rate of 1 kHz. The EMG signals underwent high-pass filtering at 100 Hz and were then integrated using LabChart software. The fold change in integrated EMG amplitude was determined by dividing the peak integrated EMG amplitude recorded during laser stimulation by the integrated EMG amplitude measured immediately prior to stimulation. Laser light at 473 nm was delivered as previously described, utilizing the following parameters: 3-second light pulses (473 nm) at frequencies of 5, 10, 20, and 40 Hz.

### Statistics and data presentation

Statistical analyses were performed using GraphPad Prism (GraphPad Software, La Jolla, CA; version 8.0). All statistical tests were two-tailed, and the specific test used for each analysis is indicated in the corresponding figure legends. For comparisons between independent groups, the Mann–Whitney U test was applied; matched or within-subject comparisons were analyzed using the Wilcoxon signed-rank test. Multifactorial datasets were evaluated using two-way analysis of variance followed by Bonferroni post hoc corrections, and paired *t* tests were used where appropriate. Statistical significance was defined as *P* < 0.05.

Behavioral experiments were conducted under computer-controlled conditions, and data acquisition and analysis were performed in an automated and unbiased manner. Animals were excluded from analysis only if post hoc histological verification revealed that viral expression, optical fiber placement, or GRIN lens positioning was outside the intended target region. No additional animals or data points were excluded. Unless otherwise stated, data are presented as mean ± SEM.

## Acknowledgements

We thank B. Li, Q. Sun, B. Arkarup and J. Wang for critically reading of the manuscript; We thank members of the Zhang lab for comments and suggestions during the entire process of the project; We thank P. Zhou and R. DONG for their assistance with motion correction of two-photon imaging. We thank the staff of the Laboratory of Animal Facility and Biosciences Central Research Facility at HKUST(GZ) for technical support. This work is supported by the National Natural Science Foundation of China (32371064 to X.Z.), the Guangdong Science and Technology Department “1+1+1” Joint Funding Program (to X.Z.), the Guangzhou Municipal Science and Technology Project (2025A04J7035 to X.Z.), the Department of Education of Guangdong Province (2025KCXTD056 to X.Z.), the Guangzhou-HKUST(GZ) Joint Funding Program (2023A03J0620 to X.Z.), and HKUST(GZ) startup Fund (to X.Z.).

## Author contributions

X.Z., Q.L. and R.X. conceptualized the study; Q.L. and R.X. and W.H. performed the Experiments; Q.L., R.X., and Y.Q. did the data analysis; Q.L.,Y.Q. and X.Z. prepared the figures; X.Z., Q.L. and Y.Q wrote the manuscript with input from all other authors.

## Data availability

Single-nucleus transcriptomic data have been deposited at GSA: https://ngdc.cncb.ac.cn/gsa/search?searchTerm=CRA035580 and available as of the date of publication. The accession number is listed in the key resources table. Other data that support the findings of this study are available from the corresponding author upon request.

## Table

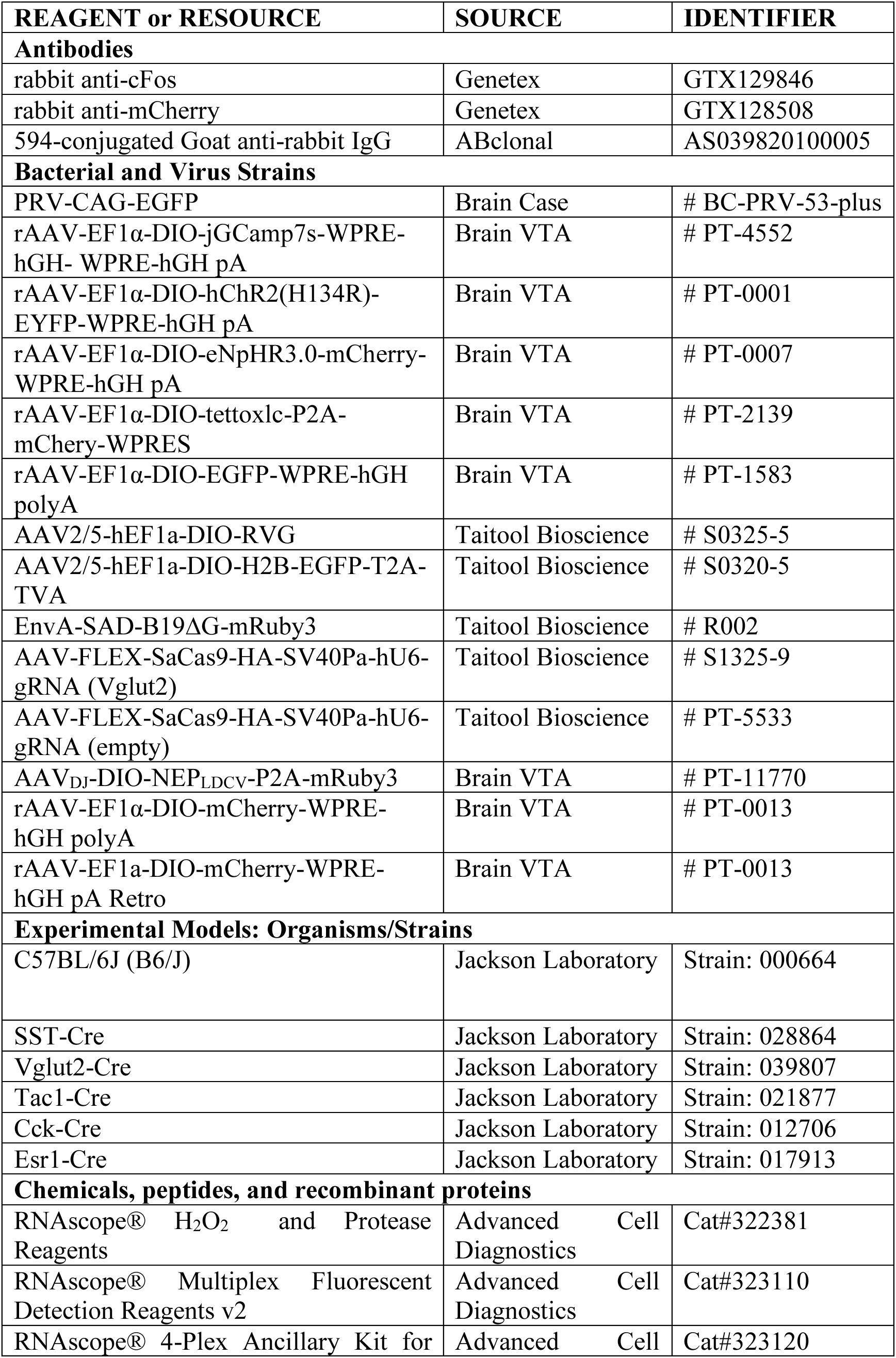
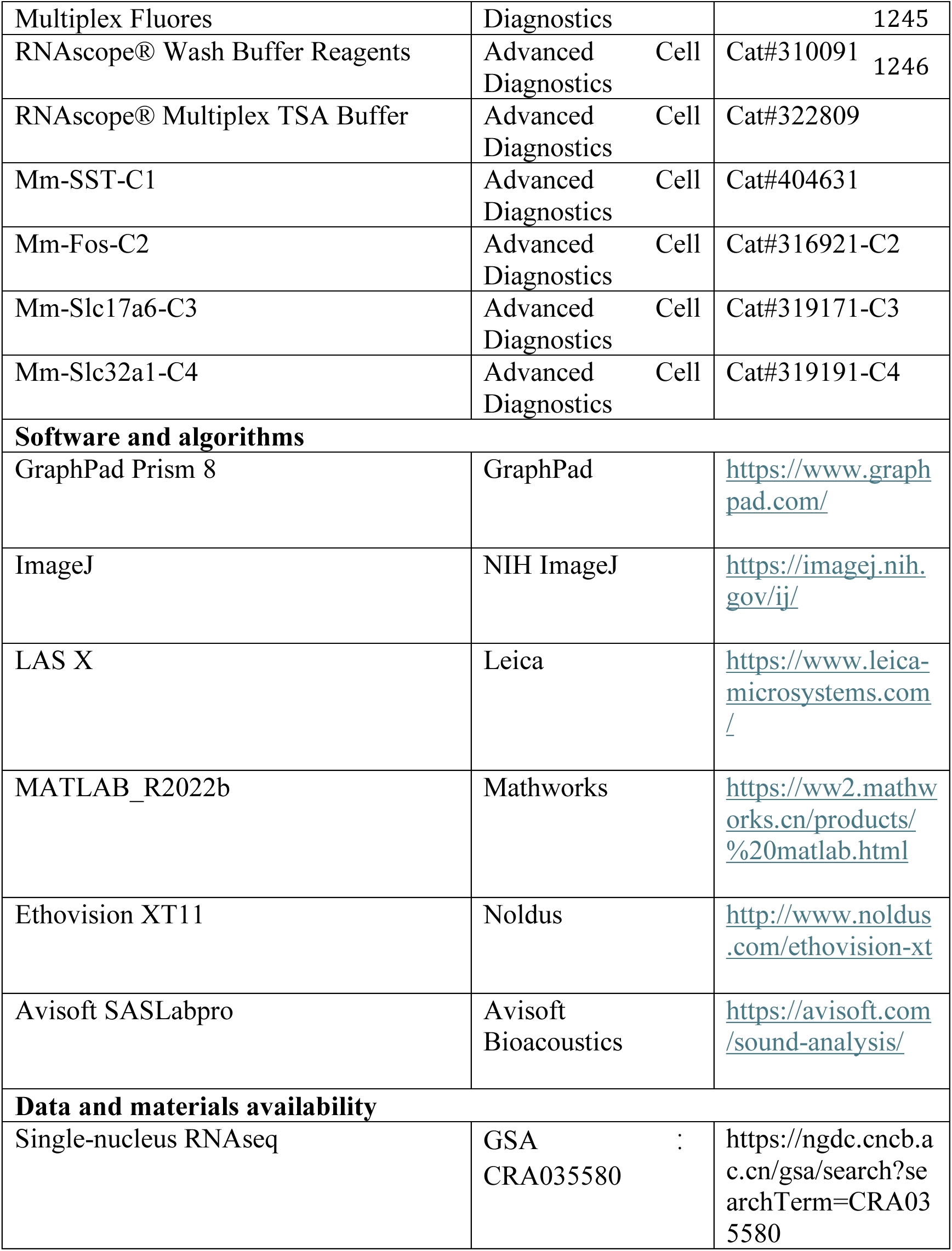

