## Supplementary Figures for "A genetically defined midbrain-pontine circuit gates vocal communication"

**Extended Data Figures:**

**Extended Data Figure 1.**

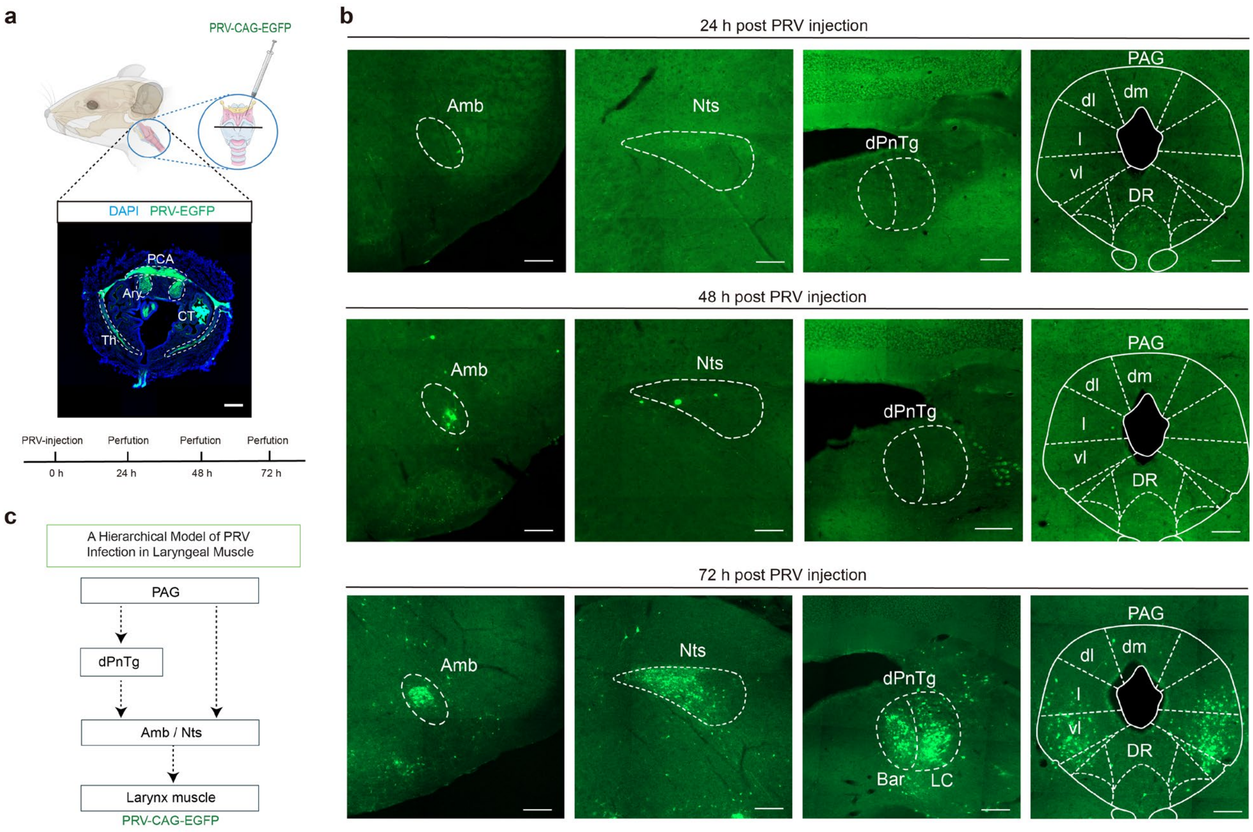


**Extended Data Fig. 1 | Time-dependent distribution of PRV-labeled neurons following PRV-EGFP injection into the laryngeal muscle. a**, Top: Schematic illustrating PRV-EGFP (PRV531) delivery into the laryngeal muscle. Bottom: Representative fluorescence image showing EGFP expression at the injection site. PCA, posterior cricoarytenoid muscle; Ary, arytenoid cartilage; CT, cricothyroid muscle; Th, thyroid cartilage. Scale bar, 300 μm. **b**, Representative coronal brainstem sections showing PRV-labeled neurons at 24, 48, and 72 h after laryngeal muscle injection. White contours delineate anatomical boundaries. At 24 hours after PRV injection, we did not observe PRV-labeled neurons in the brain. At 48 hours after injecting into the larynx, we found only sparse PRV-labeled neurons in the ambiguous nucleus (Amb) and nucleus tractus solitarius (Nts). By 72 hours post-injection, viral labeling extended to the dorsal pontine tegmentum (dPnTg), an area including Barrington's nucleus (Bar) and locus coeruleus (LC), and the lateral and ventrolateral PAG (l/vlPAG). Scale bar, 200 μm. **c**, Schematic summary of the inferred hierarchical progression of PRV trans-neuronal infection from laryngeal muscle to brainstem. PAG neurons are known to project directly to the NTS and Amb, as well as send strong projections to the dPnTg. Based on our results and existing literature, we suggest that PRV is transported retrogradely across multiple synapses, from the larynx to the l/vlPAG.

**Extended Data Figure 2.**

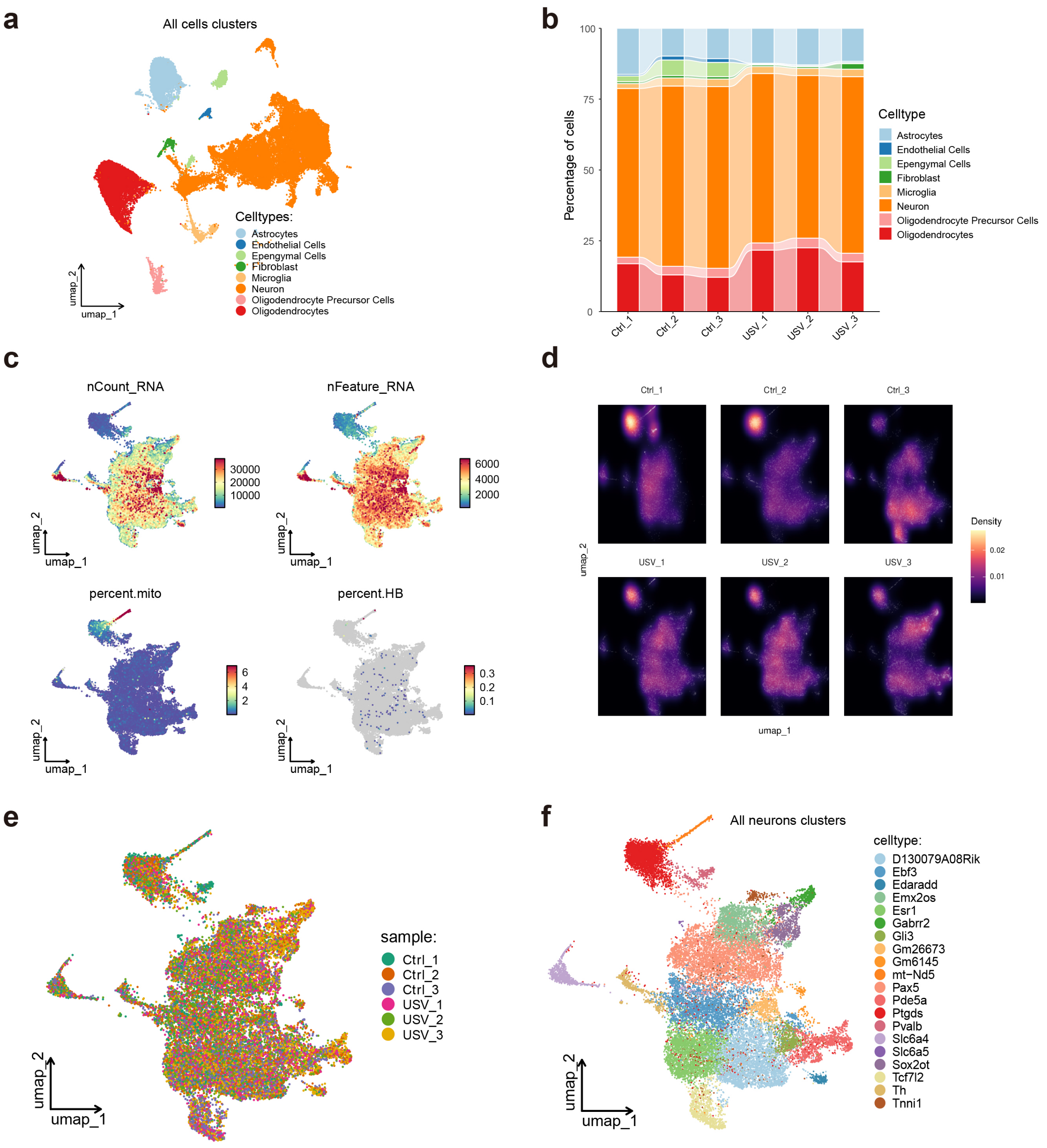


**Extended Data Fig. 2 | Cellular composition and quality control of pooled PAG single-nucleus transcriptomic datasets. a**, UMAP embedding of all captured nuclei (n = 41,054) from pooled PAG samples collected from six mice. Colors indicate major cell classes. **b**, Cell-type composition across individual samples. Each bar represents one specific sample, with colors denoting distinct cell types. **c**, UMAP projections colored by quality-control metrics, including total RNA counts (nCount_RNA), number of detected genes (nFeature_RNA), mitochondrial transcript percentage (percent.mito), and hemoglobin transcript percentage (percent.HB). **d**, Density plots showing the distribution of nuclei from individual samples in UMAP space, illustrating comparable coverage across samples. **e**, UMAP embedding colored by sample identity: Ctrl_1–3 (non-vocalizing controls) and USV_1–3 (vocalizing males). **f,** UMAP embedding of neuronal nuclei annotated into transcriptionally distinct neuronal subclusters based on marker gene expression.

**Extended Data Figure 3.**

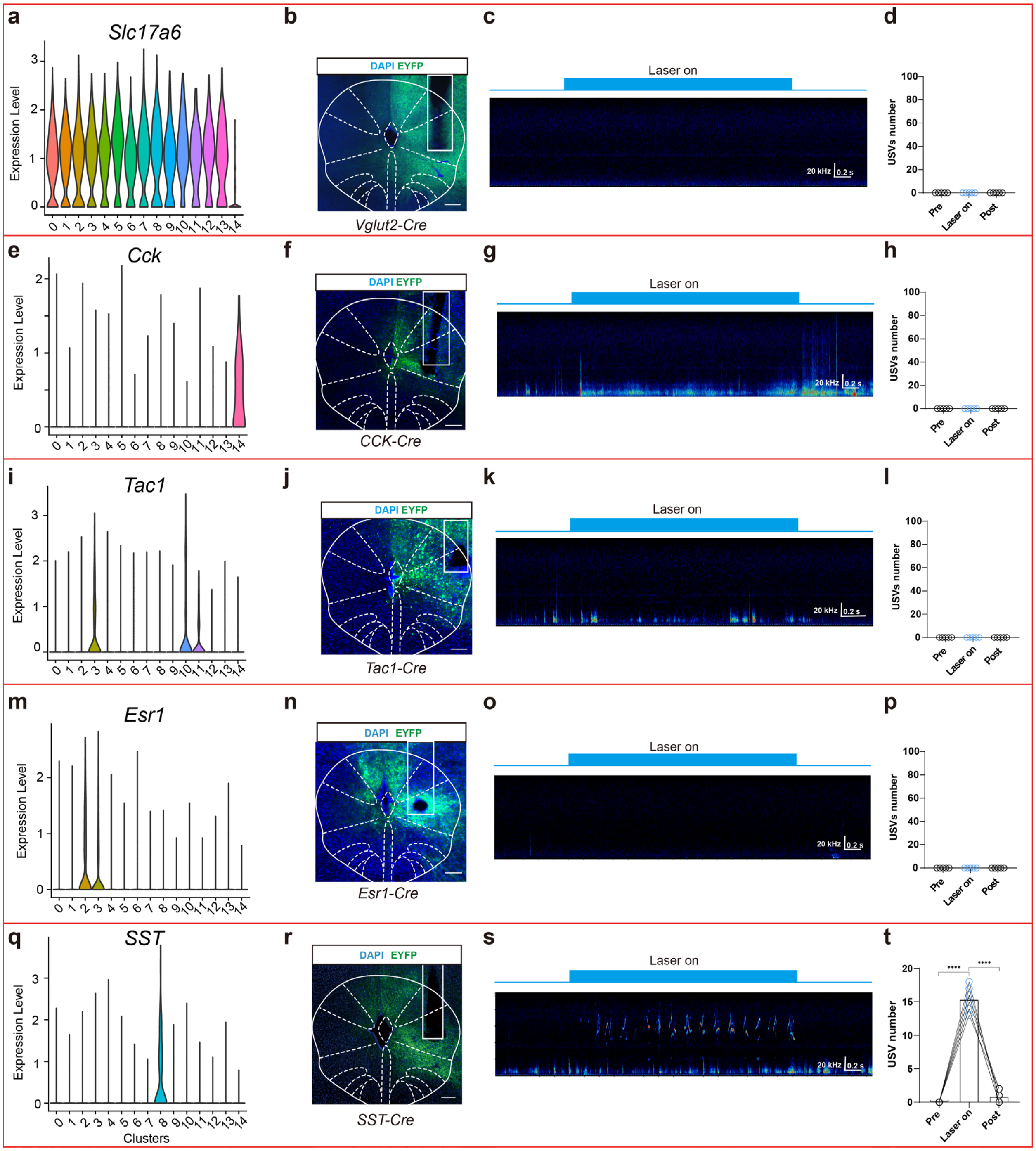


**Extended Data Fig. 3 | Functional screening of transcriptionally defined l/vlPAG neuronal subtypes identifies SST neurons as selectively sufficient to drive vocalization. a, e, i, m, q,** Violin plots showing expression levels of candidate marker genes: *Slc17a6* (general glutamatergic), *Cck* (class *I*), *Tac1* (class *II*), *Esr1* (class *III*), and *SST* (class *IV*), across l/vlPAG neuronal clusters identified by snRNA-seq. **b, f, j, n, r**, Representative coronal sections showing ChR2–EYFP expression in the l/vlPAG of *Vglut2-Cre, Tac1-Cre, Cck-Cre, Esr1-Cre,* and *SST-Cre* mice, respectively. Scale bar, 200 µm. **c, g, k, o, s**, Representative USV sonograms recorded during optogenetic stimulation (20 Hz; blue bar) of the indicated l/vlPAG neuronal populations in freely moving mice. **d, h, l, p, t**, Quantification of USV production during pre-stimulation, laser-on, and post-stimulation epochs for activation of *Vglut2* (n = 3), *Tac1* (n = 3), *Cck* (n = 3), *Esr1* (n = 3), and *SST* (n = 6) neuronal populations. Only activation of l/vlPAG$^{SST}$ neurons robustly elicited USVs. Data are mean ± s.e.m.; one-way ANOVA; ****$P < 0.0001$.

**Extended Data Figure 4.**

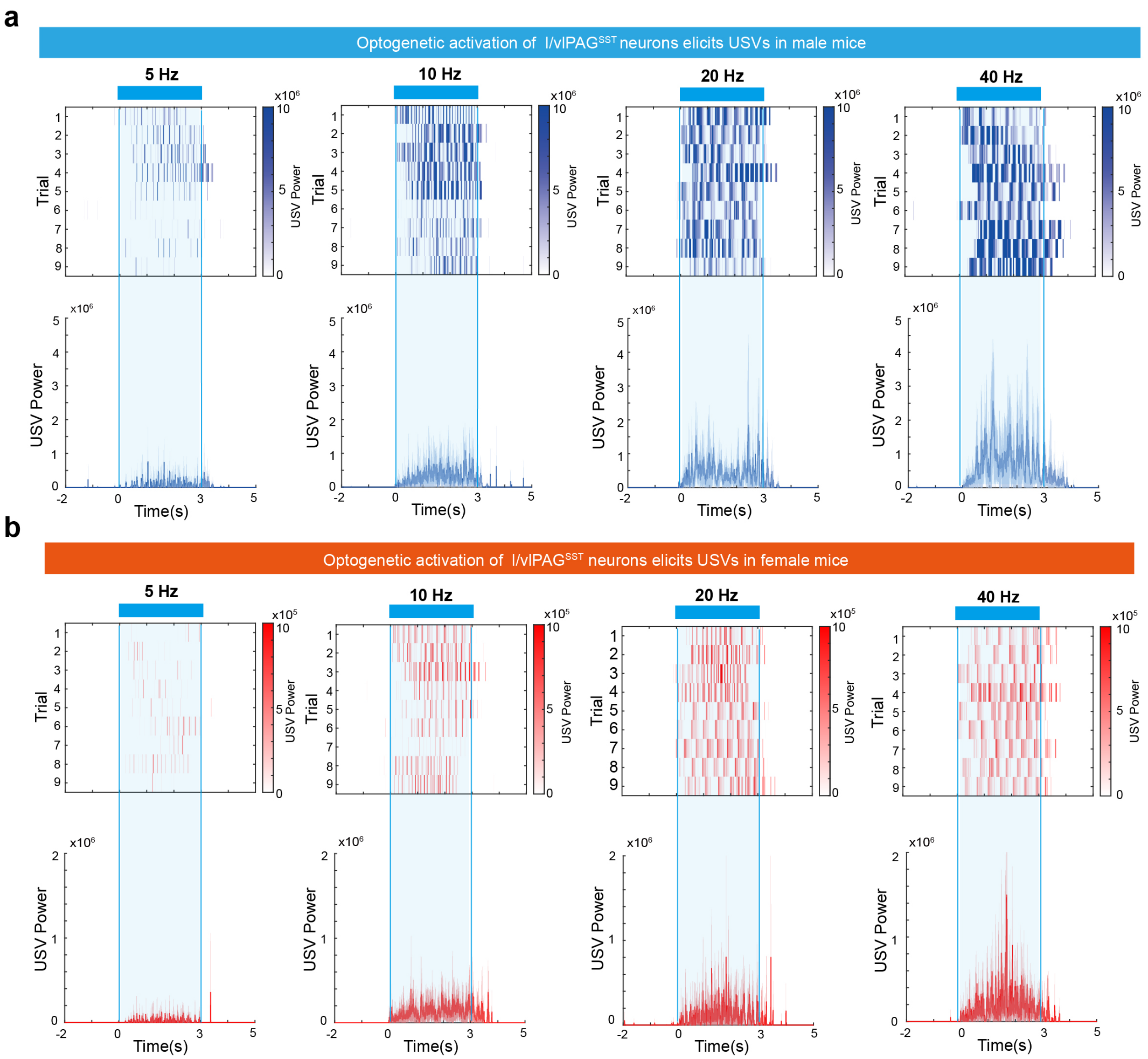


**Extended Data Fig. 4 | Optogenetic activation of l/vlPAG$^{SST}$ neurons elicits USVs in both sexes. a**, Male mice. Top, raster plots of USV syllables aligned to laser onset during optogenetic stimulation of l/vlPAG$^{SST}$ neurons at 5, 10, 20, and 40 Hz. Bottom, corresponding trial-averaged USV power traces, with the stimulation window indicated by shading. **b,** Female mice. Top, USV syllable rasters aligned to laser onset at the same stimulation frequencies. Bottom, corresponding mean USV power traces. Across both sexes, increasing stimulation frequency produced progressively stronger and more sustained USV output.

**Extended Data Figure 5.**

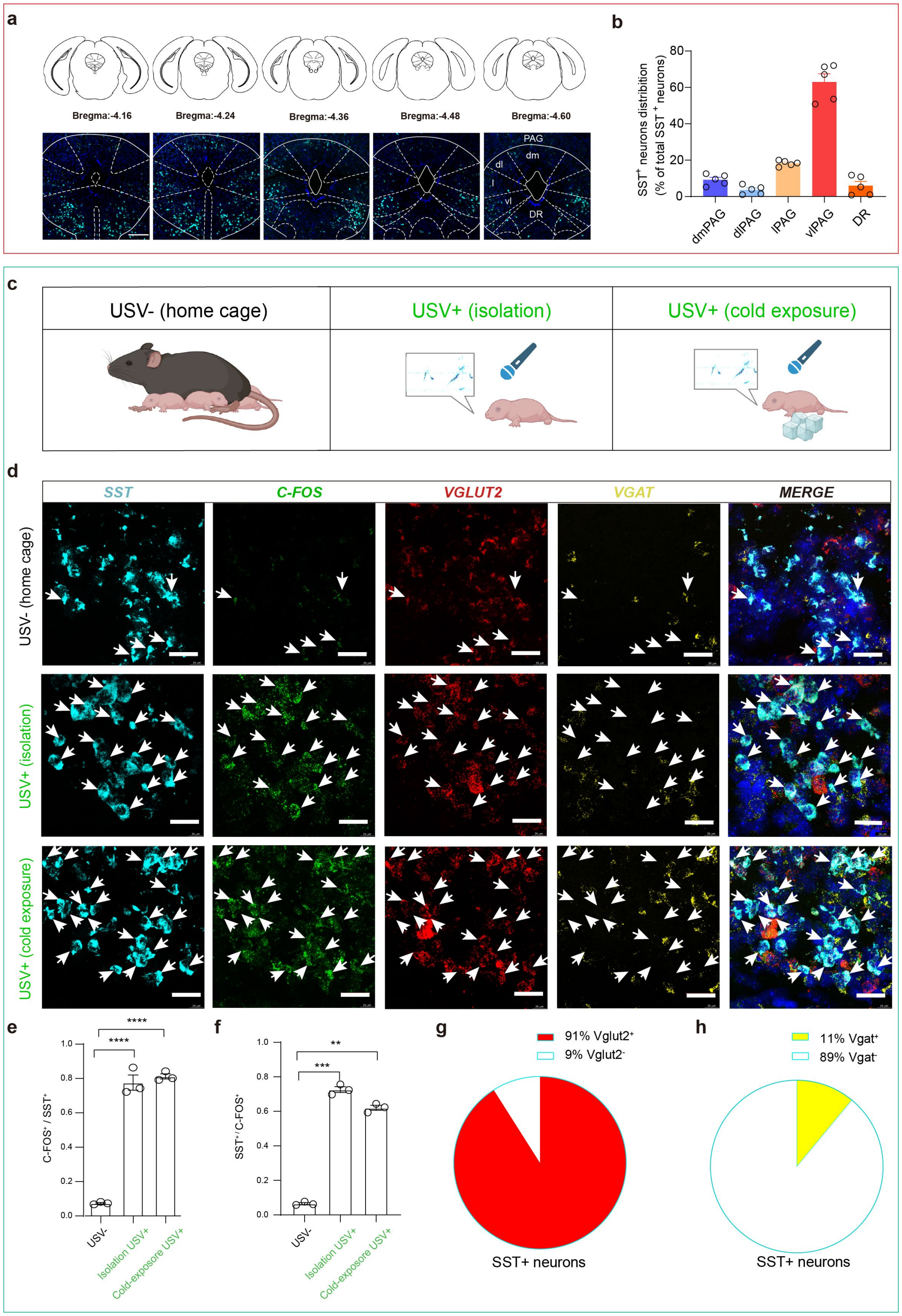


**Extended Data Fig. 5 | Spatial organization of *SST*-expressing neurons along the rostrocaudal axis of the adult PAG and their activation during pup ultrasonic vocalizations.**

**a**, Representative RNAscope images from adult male mice showing the spatial distribution of $SST^{+}$ neurons across rostrocaudal axis of the PAG (Bregma −4.16 to −4.60 mm). Upper panels show corresponding coronal atlas sections; lower panels show RNAscope detection of *SST* mRNA (cyan) overlaid on *DAPI*. Dashed outlines denote PAG subregions. Scale bar, 200 µm. **b**, Quantification of the proportion of $SST^{+}$ neurons across PAG subregions, including dorsomedial (dmPAG), dorsolateral (dlPAG), lateral (lPAG), ventrolateral (vlPAG), and dorsal raphe (DR). Data are expressed as percentage of total $SST^{+}$ neurons within the PAG (mean ± s.e.m.). **c**, Schematic of experimental conditions used to induce pup USVs: USV- home-cage control, and USV+ conditions induced by isolation or cold exposure for *Fos* mRNA detection. **d**, Representative RNAscope images comparing *SST*, *Fos*, *Vglut2*, and *Vgat* expression in l/vlPAG between USV- and USV+ pups. Arrowheads denote l/vlPAG$^{SST}$ neurons. Scale bar, 50 µm. **e**, Percentage of *SST* neurons expressing *Fos* under USV+ versus USV- conditions (n = 3 pups/group; unpaired t-test, ****$P$ < 0.0001). **f**, Proportion of l/vlPAG$^{Fos}$ neurons that express *SST* in vocalizing pups (n = 3 pups/group; unpaired t-test, **$P$ < 0.01, ***$P$ < 0.001). **g**, Fraction of l/vlPAG$^{SST}$ neurons co-expressing *Vglut2* ($n$ = 6 pups). **h**, Fraction of l/vlPAG$^{SST}$ neurons co-expressing *Vgat* ($n$ = 6 pups).

**Extended Data Figure 6.**

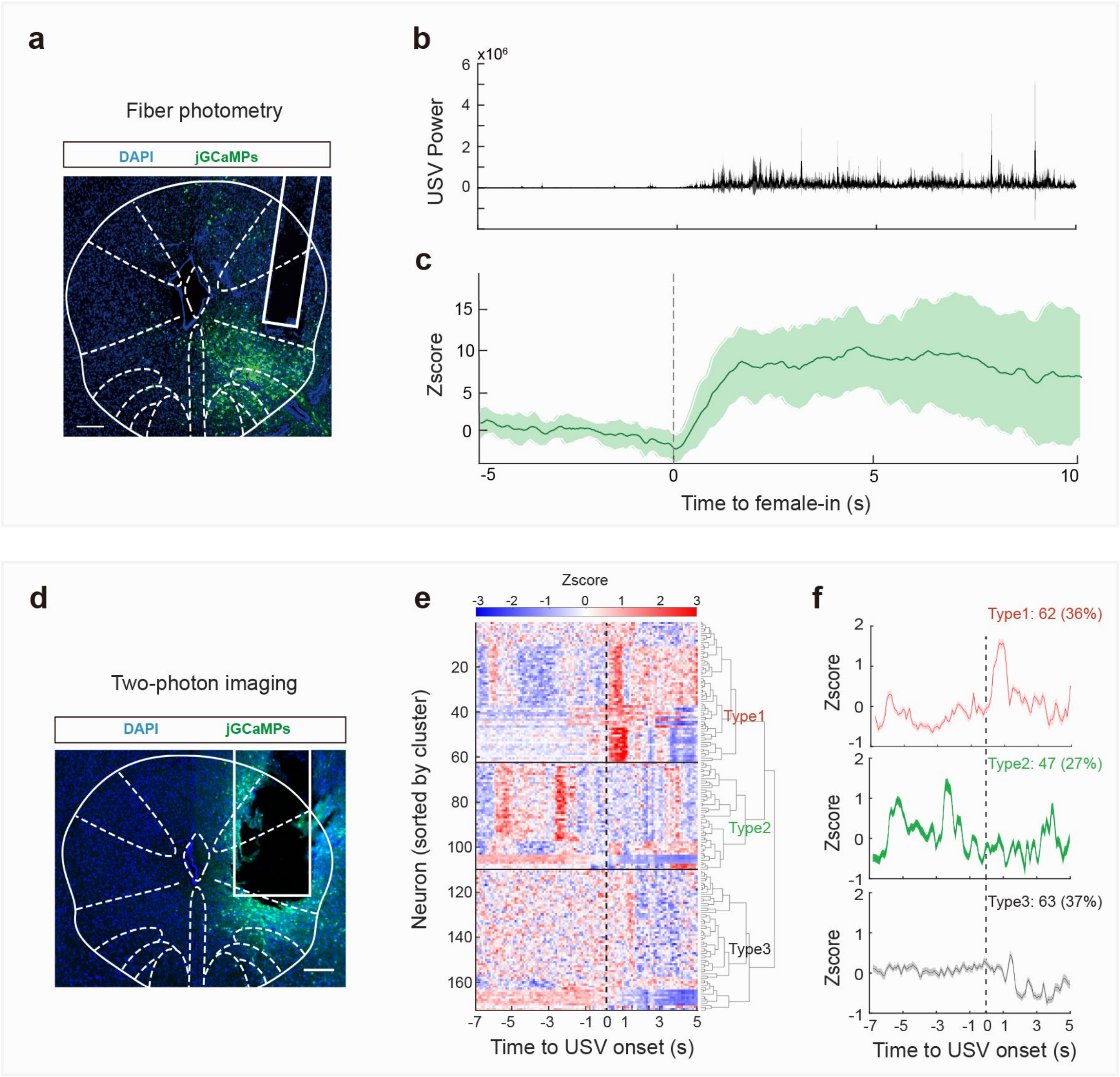


**Extended Data Fig. 6 | l/vlPAG$^{SST}$ neuronal activity correlates with USV generation at both population and single-cell levels. a**, Coronal brain image showing jGCaMP7s expression and optic fiber trajectory in *SST-Cre* mice. Scale bar, 200 μm. **b**, Averaged USV power recorded from all fiber photometry male mice during courtship interaction. **c**, Averaged calcium signal (z-score) aligned to introduction of female (n = 8 mice). Shaded area represents 95% confidence interval. **d**, Confocal image showing jGCaMPs expression and GRIN lens trajectory above l/vlPAG for freely moving two-photon imaging. Scale bar, 200 μm. **e**, Heat map of single-cell calcium activity aligned to USV onset, showing hierarchical clustering of all recorded neurons (n = 172 neurons, 5 mice). **f**, Mean activity traces for each cluster: Type 1 (62 cells, 36%), Type 2 (47 cells, 27%), and Type 3 (63 cells, 37%). Shaded areas represent SEM.

**Extended Data Figure 7.**

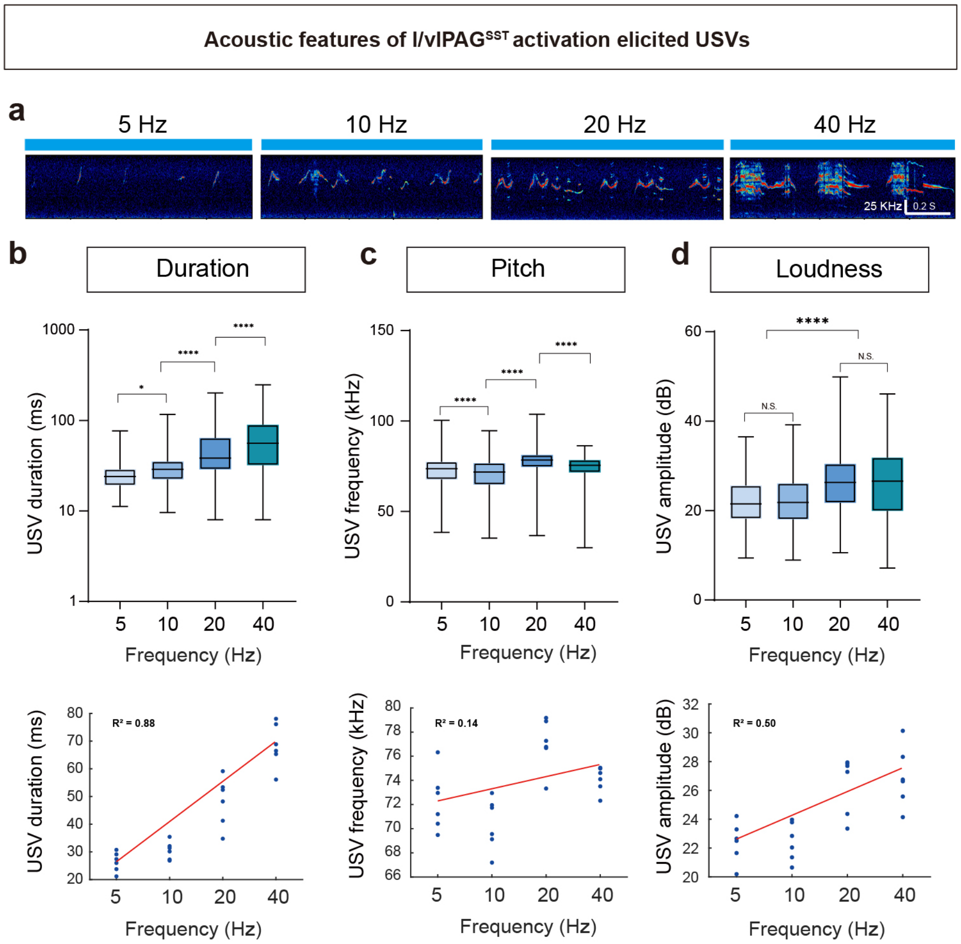


**Extended Data Fig. 7 | Optogenetic activation of l/vlPAG$^{SST}$ neurons scales USV duration.** **a**, Representative sonograms of optogenetically evoked USVs at stimulation frequencies of 5, 10, 20, and 40 Hz. **b–d,** Quantification of USV acoustic features during frequency-dependent stimulation: (**b**) syllable duration, (**c**) pitch, and (**d**) loudness (n = 6 mice). Box plots show minimum, maximum, and interquartile range. Statistical comparisons were performed using one-way ANOVA (*****p < 0.0001, *p < 0.05, ns = not significant*). Bottom, linear regression showing frequency-dependent scaling of USV duration ($R^2 = 0.88$), with weaker relationships for pitch ($R^2 = 0.14$) and amplitude ($R^2 = 0.50$).

**Extended Data Figure 8.**

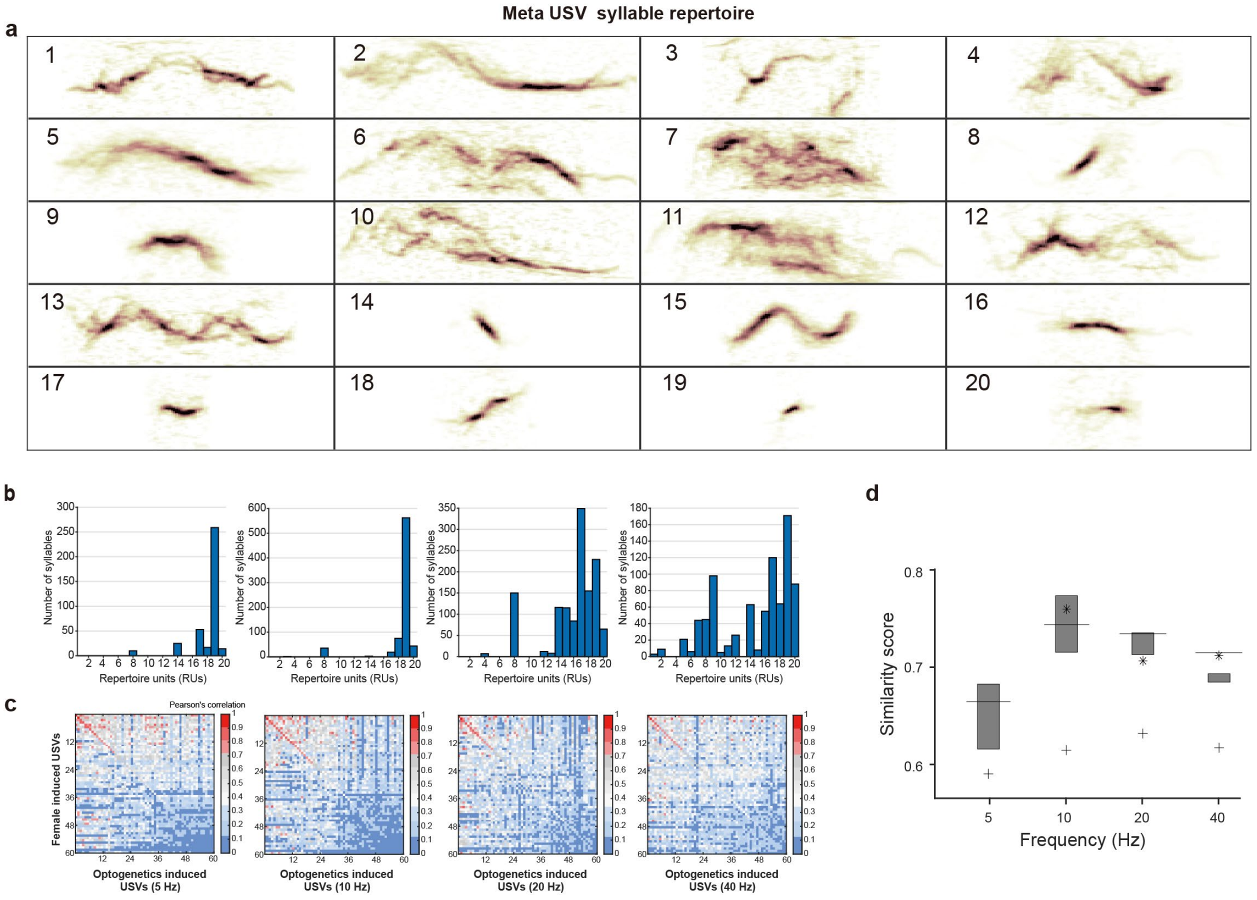


**Extended Data Fig. 8 | Frequency-dependent activation of l/vlPAG$^{SST}$ neurons increases USV syllabic diversity and the similarity to natural courtship vocalizations. a**, MUPET conducts a cluster analysis of repertoire unit types (RUs) to create a master repertoire of RUs clusters. **b**, Distribution of RU usage across USVs elicited by optogenetic stimulation at 5, 10, 20, and 40 Hz, demonstrating frequency-dependent changes in syllable composition. **c**, Pearson's correlation matrices comparing USVs evoked at each stimulation frequency to natural USVs recorded from wild-type males interacting with conspecific females, revealing broad and diverse syllable structures across stimulation conditions. Among all tested frequencies, 10-Hz stimulation produced USVs with the highest similarity to natural courtship vocalizations. **d**, Quantification of similarity scores between natural USVs and optogenetically evoked USVs across stimulation frequencies (5, 10, 20, and 40 Hz) in ChR2-expressing l/vlPAG$^{SST}$ mice (n = 6). The asterisk (∗) denotes the Pearson correlations for the top 5% of the most frequently used repertoire units, where the boxplot shows the mean and interquartile range of these correlations, and the plus sign (+) shows the correlation of the top 95% of the most frequency used repertoire units.

**Extended Data Figure 9.**

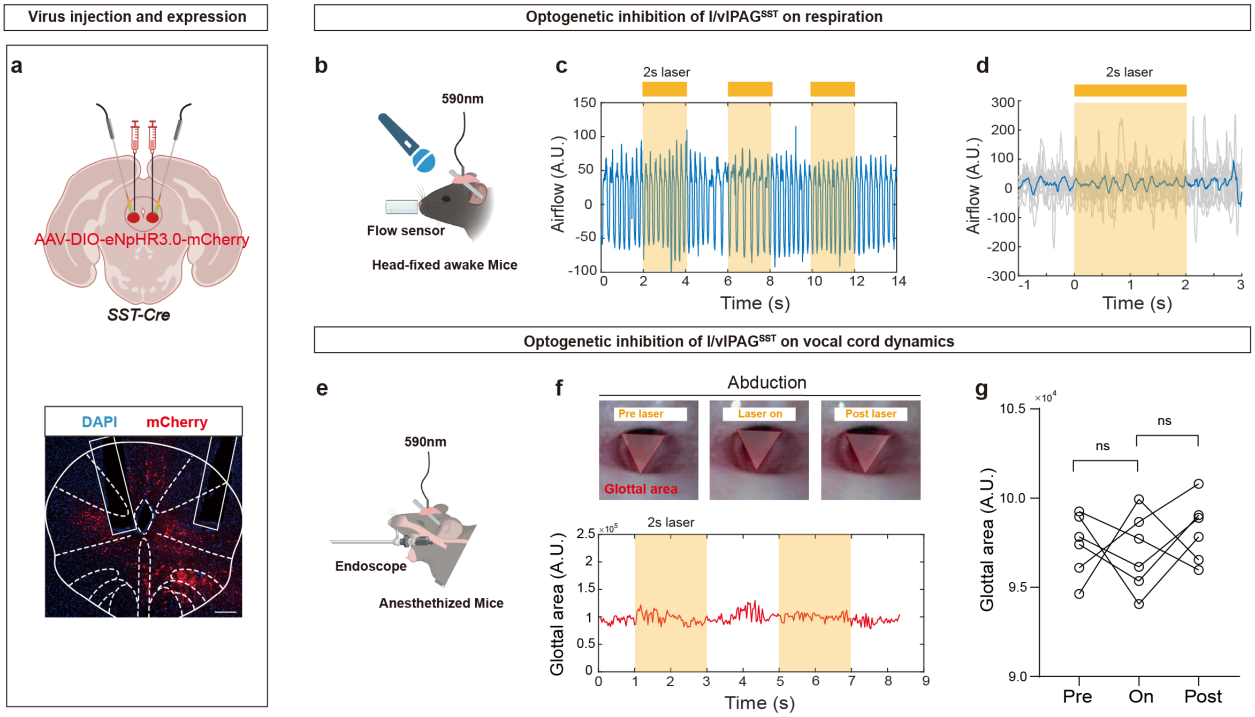


**Extended Data Fig. 9 | Optogenetic inhibition of l/vlPAG$^{SST}$ neurons does not alter respiratory patterns or vocal fold dynamics. a**, Top, schematic illustrating bilateral injection of AAV-EF1α-DIO-eNpHR3.0-mCherry into the l/vlPAG and optical fiber implantation in *SST-Cre* mice. Bottom, fluorescence image showing eNpHR3.0-mCherry bilateral expression in l/vlPAG neurons and the optical fiber trajectory. Scale bar, 200 μm. **b**, Schematic diagram of the head-fixed respiration-monitoring setup in awake mice. **c**, Representative and **d**, average airflow traces recorded during 2-s optogenetic inhibition of l/vlPAG$^{SST}$ neurons. Twelve trials are aligned to the laser onsets and overlaid; yellow shading indicates laser duration. **e**, Schematic of laryngeal endoscopy setup to visualize vocal fold movement under anesthesia. **f**, Top, endoscopic images of glottal area changes before, during, and after optogenetic inhibition. Bottom, continuous measurement of glottal area aligned to the laser period (yellow shading). **g**, Quantification of glottal area across pre-, during-, and post-laser inhibition (n = 6 mice). No significant differences were detected (paired t-test; ns, $p \geq 0.05$).

**Extended Data Figure 10.**

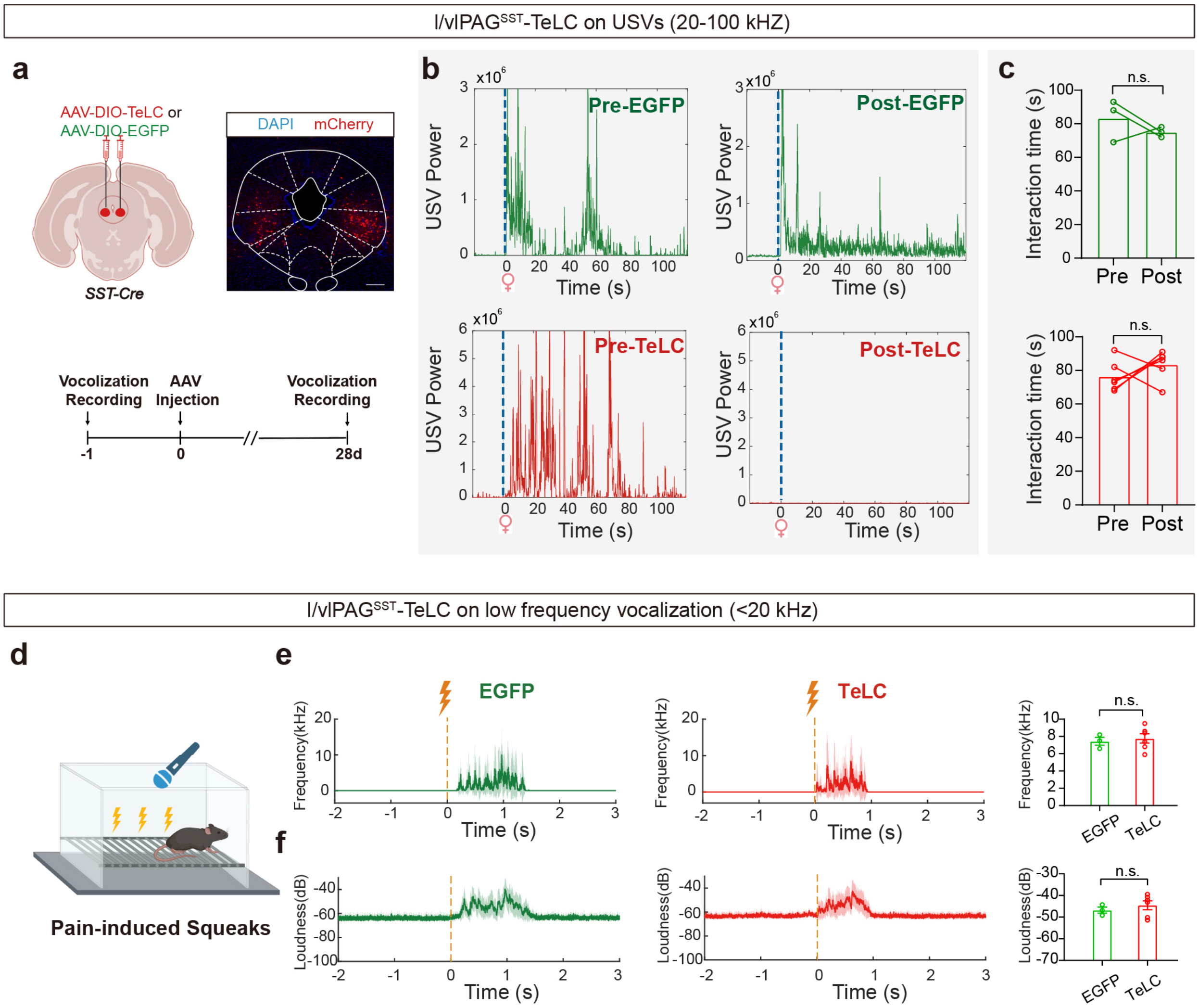


**Extended Data Fig. 10 | Synaptic silencing of l/vlPAG$^{SST}$ neurons abolishes courtship USVs, but does not affect low-frequency vocalizations.** **a**, Top: schematic showing the bilateral injection of the AAV-EF1α-DIO-TeLC-mCheery or AAV-EF1α-DIO-EGFP virus into the l/vlPAG of *SST-Cre* mice. Right, representative image of TeLC-mCherry expression in the l/vlPAG. Scale bar, 200 μm. Bottom, experimental timeline: male–female interaction-induced USVs were recorded 1 day prior to virus injection, followed by bilateral viral delivery at day 0, and repeated USV recordings 28 days later after stable TeLC expression and synaptic blockade. **b**, Representative USV power traces from *SST-Cre* mice expressing EGFP (green) or TeLC (red) during female-evoked courtship sessions before (Pre) and 4 weeks after (Post) viral injection. Blue dashed line denotes female introduction. **c**, Quantification of interaction duration during the 3-min social assay (EGFP, n = 3; TeLC, n = 6). ns, $p \geq 0.05$; paired t-test. **d**, Schematic of footshock-evoked audible squeak recording. **e, f**, Frequency (**e**) and amplitude (**f**) traces of pain-evoked squeaks in EGFP-expressing (green) and TeLC-expressing (red) mice. Yellow dashed line marks footshock onset. No significant differences were detected (ns, $p \geq 0.05$).

**Extended Data Figure 11.**

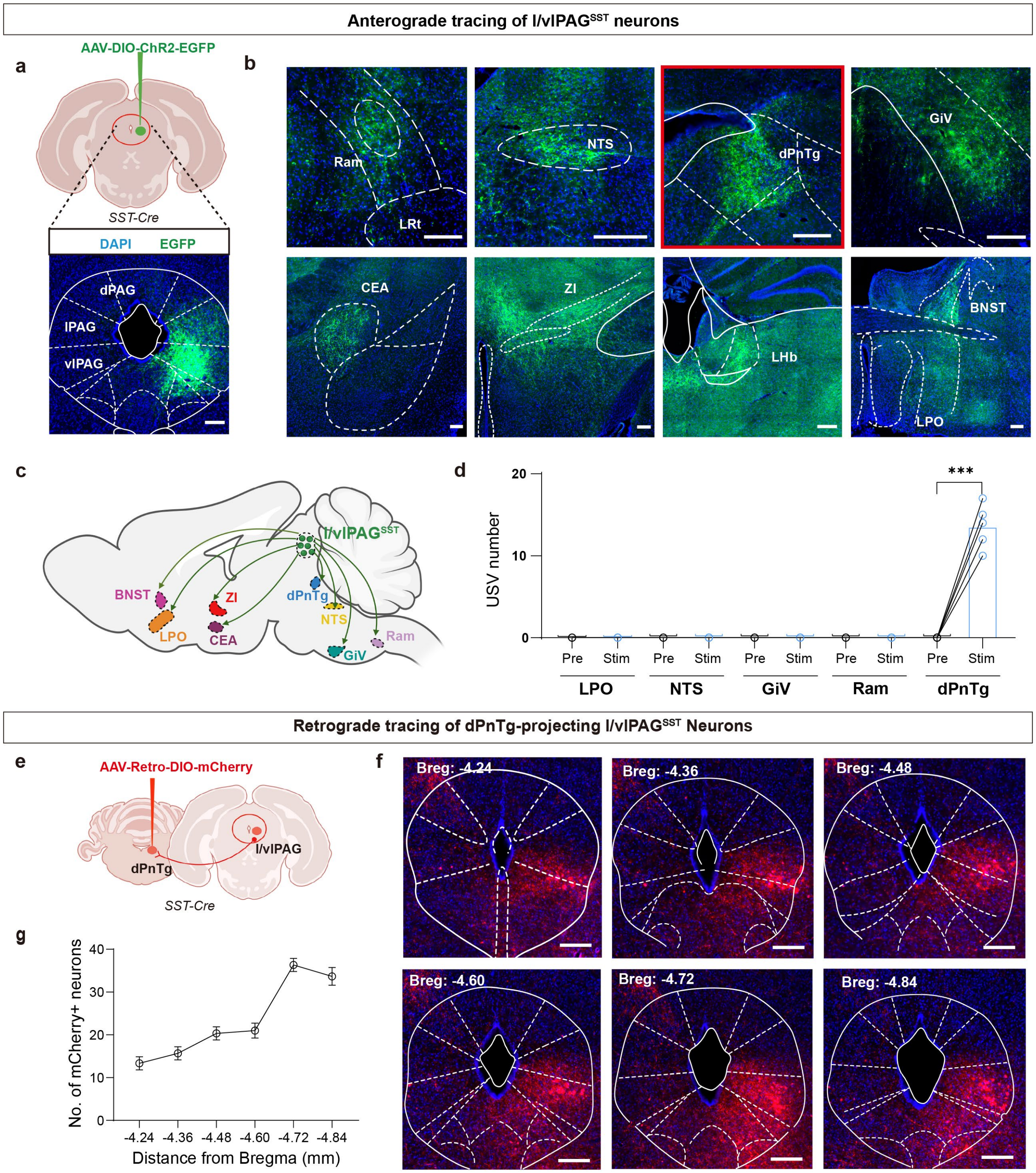


**Extended Data Fig. 11 | Anterograde and functional circuit mapping of l/vlPAG$^{SST}$ neurons.** **a**, Schematic of AAV-DIO-ChR2-EGFP delivery into the unilateral l/vlPAG of *SST-Cre* mice (top) and representative expression in l/vlPAG$^{SST}$ neurons (bottom). Scale bar, 100 μm. **b**, Representative images of downstream terminal arising from l/vlPAG$^{SST}$ neurons across multiple brainstem and forebrain areas. Scale bar, 200 μm; n = 3 mice. **c**, Diagram summarizing major projection sites of l/vlPAG$^{SST}$ neurons, including LPO, BNST, ZI, CeA, GiV, NTS, and dPnTg. **d**, Quantification of USVs evoked by selective optogenetic activation of l/vlPAG$^{SST}$ axon terminals within individual downstream regions (LPO: n = 3; NTS: n = 3; GiV: n = 3; Ram: n = 3; dPnTg: n = 5). Robust USV production was observed only following dPnTg terminal stimulation (***$p < 0.001$, paired t-test). **e**, Schematic of AAV-Retro-DIO-mCherry injection into the unilateral dPnTg of *SST-Cre* mice to identify dPnTg-projecting l/vlPAG$^{SST}$ neurons (n = 3). **f**, Representative serial coronal sections showing mCherry-labeled dPnTg-projecting neurons distributed throughout the l/vlPAG. Scale bar, 200 μm. **g**, Quantification of mCherry-positive neurons along the rostro-caudal l/vlPAG axis (Bregma −4.24 to −4.84 mm, n = 3 mice). Ram: retroambiguus nucleus; Sol: nucleus of the solitary tract; dPnTg: dorsal pontine tegmentum; GiV: gigantocellular reticular nucleus; CEA: central amygdaloid nucleus; ZI: zona incerta; LHb: lateral habenular nucleus; BNST: bed nucleus of the stria terminalis; LPOA: lateral preoptic area.

**Extended Data Figure 12.**

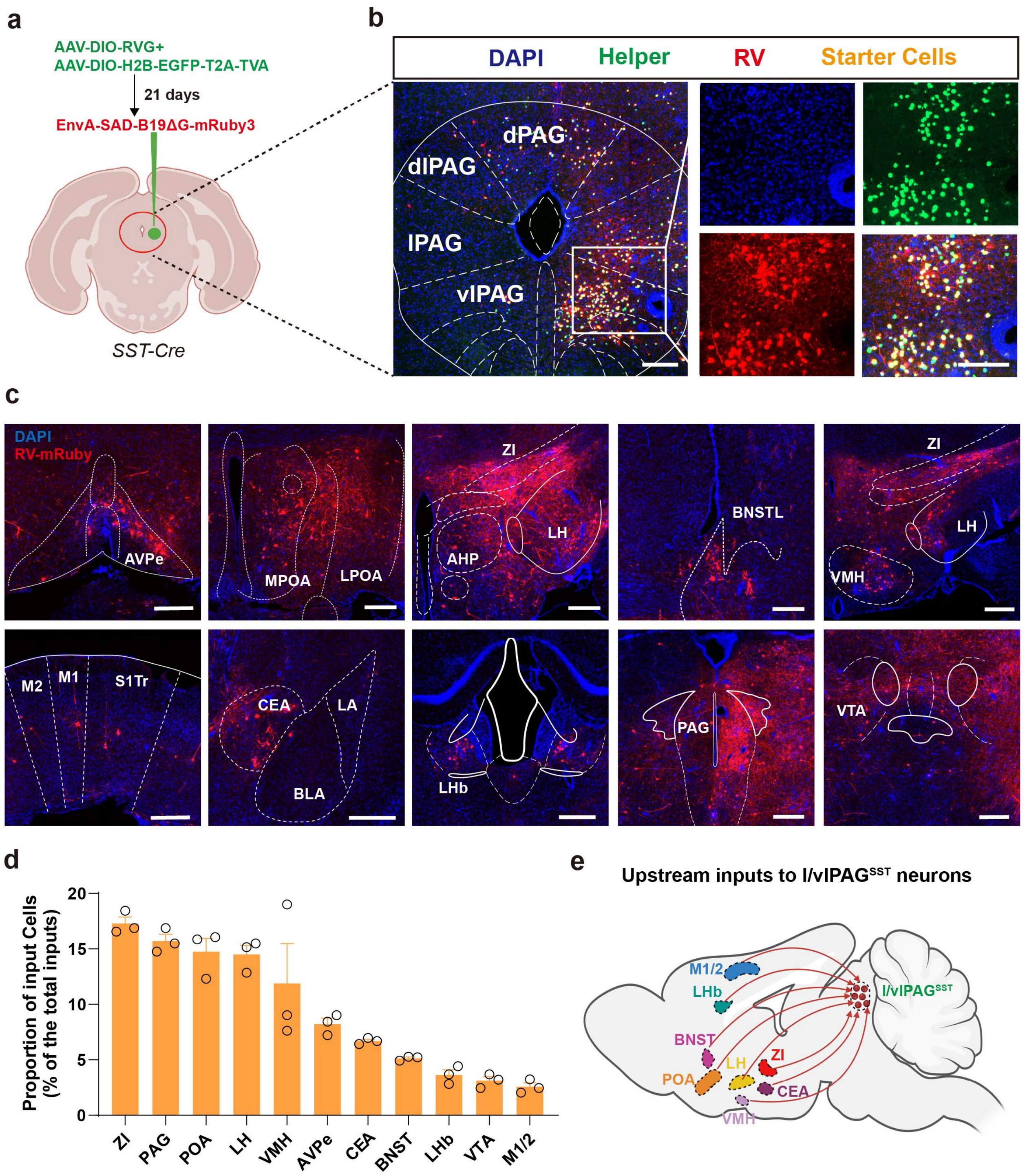


**Extended Data Fig. 12 | Monosynaptic retrograde tracing identifies distributed forebrain inputs to l/vlPAG$^{SST}$ neurons. a**, Schematic of the monosynaptic rabies tracing strategy. *SST-Cre* mice received l/vlPAG injections of Cre-dependent helper AAVs expressing rabies glycoprotein (RG) and the TVA receptor fused to EGFP. Twenty-one days later, EnvA-pseudotyped, glycoprotein-deleted rabies virus expressing mRuby3 was injected into the same site to restrict transsynaptic labeling to monosynaptically connected presynaptic neurons. **b**, Confocal images of the injection site showing helper-expressing cells (green), rabies-infected cells (red), and starter cells (yellow) confined to the l/vlPAG. Scale bar, 100 μm. **c**, Representative images of major forebrain and midbrain input regions labeled by rabies tracing, including hypothalamic, amygdalar, extended amygdala, and basal forebrain nuclei, as well as cortical and midbrain sites. Scale bars, 200 μm. **d**, Quantification of the proportional distribution of monosynaptic inputs to l/vlPAG$^{SST}$ neurons across brain regions (mean ± s.e.m.; individual mice shown as circles). **e**, Schematic summary of upstream inputs to l/vlPAG$^{SST}$ neurons, highlighting convergent input from emotional and motivational forebrain regions. AVPe: anteroventral periventricular nucleus; M/LPOA: medial/lateral preoptic area; ZI: zona incerta; LH: lateral habenular nucleus; BNST: bed nucleus of the stria terminalis; VMH: ventromedial hypothalamic nucleus; M1: primary motor cortex; M2: secondary motor cortex; CEA: central amygdaloid nucleus; LHb: lateral habenular nucleus; VTA: ventral tegmental area.

**Extended Data Figure 13.**

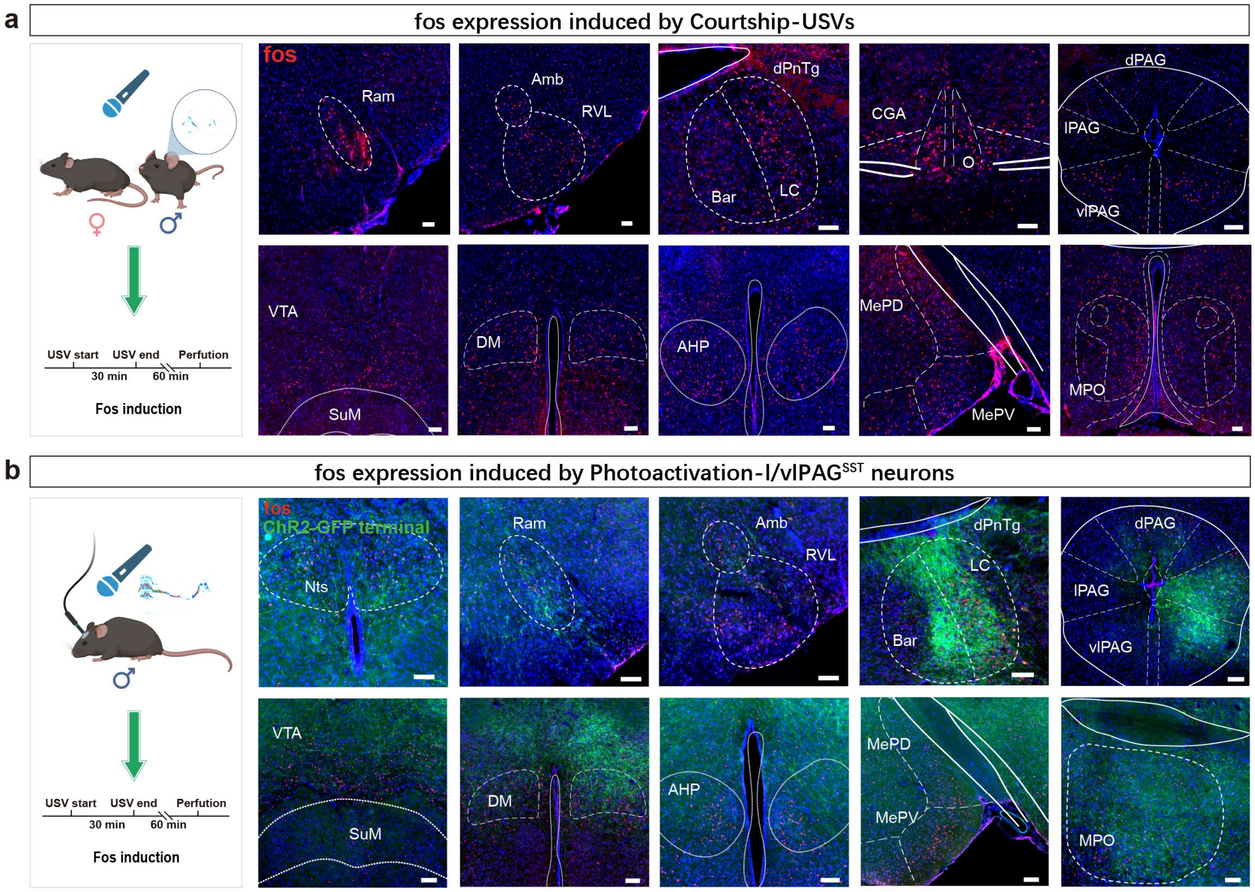


c

Courtship-USVs V.S. Photoactivation-USVs

| Brain region | Nts | Ram | Amb | RVL | dPnTg | CGA | l/vlPAG | VTA | DM | AHP | Mea | MPO |
|---|---|---|---|---|---|---|---|---|---|---|---|---|
| C-fos (courtship USVs) | - | + | + | + | + | + | + | + | + | + | + | + |
| C-fos (photoactivated USVs) | + | + | + | + | + | - | + | + | + | + | + | + |
| ChR2 Density | * | * | * | * | *** | - | **** | - | - | - | - | ** |

**Extended Data Fig. 13 | Brain-wide Fos activation associated with courtship and optogenetically evoked USVs. a**, Left, schematic of the behavioral paradigm used to induce Fos expression during male courtship vocalizations. Male mice were exposed to females to elicit USVs, followed by perfusion after the indicated interval. Right, representative images showing C-fos expression across multiple brainstem and forebrain regions in male mice emitting courtship USVs. Fos-positive neurons were observed in regions including the brainstem, midbrain, and hypothalamus. Scale bar, 100 μm; $n$ = 3 mice. **b**, Left, schematic of optogenetic activation of l/vlPAG$^{SST}$ neurons to induce USVs and Fos expression. Right, representative images of C-fos expression across corresponding brain regions following l/vlPAG$^{SST}$ photoactivation–evoked vocalizations. ChR2-GFP labeled axon terminals are also shown in the corresponding brain regions. Scale bar, 100 µm; *n* = 3 mice. **c**, Summary table comparing brain regions exhibiting Fos activation during courtship-induced versus photoactivation-induced USVs. "+" indicates detectable Fos induction and "–" indicates no detectable activation. The relative density of ChR2-positive terminals within each region is indicated on a qualitative scale (**** strong, *** moderate, ** weak, * sparse, – absent). Nts: Nucleus of the solitary tract; Ram: Retroambiguus nucleus;Amb: Nucleus ambiguous; RVL: Rostral ventrolateral medulla; dPnTg: Dorsal pontine tegmentum; Bar: Barrington's nucleus; LC: Locus coeruleus; CGA: Central gray of the pons;dPAG: Dorsal periaqueductal gray; lPAG: Lateral periaqueductal gray; vlPAG: Ventrolateral periaqueductal gray;VTA: Ventral tegmental area; SuM: Supramammillary nucleus; DM: Dorsomedial hypothalamic nucleus; AHP: Anterior hypothalamic area; MePD: Posterodorsal medial amygdala; MePV: Posteroventral medial amygdala; MeA: Medial amygdala; MPO: Medial preoptic area.

### Supplementary Videos

**Supplementary Video 1. Photoactivation of l/vlPAG$^{SST}$ neurons with different frequencies triggers USVs.**

Top: video of optogenetic stimulation of l/vlPAG$^{SST}$ neurons expressing ChR2. Bottom: Corresponding USV sonogram. The same individual mouse received 3-s light pulses delivered at 5 Hz, 10 Hz, 20 Hz, and 50 Hz. (5 Hz, 3 s laser on; 10 Hz, 3 s laser on; 20 Hz, 3 s laser on; 40 Hz, 3 s laser on)

**Supplementary Video 2. Fiber photometry recordings of l/vlPAG$^{SST}$ neuronal activity during courtship USVs (4× speed).**

Top: Video of an *SST-Cre* male mouse interacting with a female conspecific. Middle: Bulk calcium signals recorded from l/vlPAG$^{SST}$ neurons before, during, and after courtship USV production. Bottom: Corresponding USV sonogram. Video displayed at 4× real-time speed.

**Supplementary Video 3. Freely moving two-photon calcium imaging of l/vlPAG$^{SST}$ neurons during courtship USVs.**

Top left: Two-photon imaging of single-neuron activity in l/vlPAG$^{SST}$ neurons; colored contours indicate example neurons. Top right: Video of an *SST-Cre* male mouse interacting with a female conspecific. Middle: Calcium activity traces from corresponding example neurons aligned to female introduction and the onset of courtship USVs. Bottom: Corresponding USV sonogram.

**Supplementary Video 4. Photoactivation of l/vlPAG$^{SST}$ neurons induces flat-expiration airflow.**

Top left: Simultaneous USV recording and respiratory monitoring in head-fixed mice. Top right: Respiratory traces evoked by photoactivation of l/vlPAG$^{SST}$ neurons at increasing stimulation frequencies (5, 10, 20, and 40 Hz). Bottom: Corresponding USV sonogram at increasing stimulation frequencies (5, 10, 20, and 40 Hz).

**Supplementary Video 5. Photoactivation of l/vlPAG$^{SST}$ neurons drives frequency-dependent vocal-cord adduction.**

Left: Endoscopic visualization of vocal-cord movements in anesthetized mice during optogenetic stimulation. Right: Illustration of stimulation-evoked adduction duration at increasing stimulation frequencies (5, 10, 20, and 40 Hz).

**Supplementary Video 6. Photoinhibition of l/vlPAG$^{SST}$ neurons abolishes ongoing courtship USVs.**

Top: Video of optogenetic inhibition of l/vlPAG$^{SST}$ neurons (NpHR expression in *SST-Cre* male mice) during interaction with female a mouse. Bottom: USV sonogram showing the immediate disruption of ongoing courtship vocalizations during photoinhibition.

**Supplementary Video 7. Synaptic silencing of l/vlPAG$^{SST}$ neurons abolishes courtship USVs (5× speed).**

Top: Video of SST-Cre male mice interacting with a female mouse. Bottom: USV sonogram showing the complete loss of courtship vocalizations following TeLC-mediated blockade of synaptic output from l/vlPAG$^{SST}$ neurons.

**Supplementary Video 8. Photoactivation of l/vlPAG$^{SST}$ axon terminals in dPnTg evokes USVs.**

Top: Video of optogenetic stimulation of ChR2-expressing l/vlPAG$^{SST}$ axon terminals in dPnTg. Bottom: USV sonogram showing vocalizations elicited by terminal stimulation.
